# Interrogation of single-neuron functional connectivity in the cortex and hippocampus via fast cross-layer all-optical physiology

**DOI:** 10.1101/2023.08.15.553353

**Authors:** Chi Liu, Yuejun Hao, Yi Zhong, Lingjie Kong, Bo Lei

## Abstract

The interrogation of functional neural circuits is crucial for uncovering how the brain works during diverse behaviors. Multi-plane neurophysiological measurement systems with high temporal resolution are indispensable, especially for dissecting inter-layer functional connectivity. Here, we develop a cross-layer all-optical physiology system (CLAOP) that enables the simultaneous recording and manipulation of single-neuron activities in multiple neuronal layers, with axial intervals as large as 530 μm, at high temporal resolutions. Based on spatiotemporal multiplexing, our system enables all-optical analysis with a high frame rate up to 396 Hz and minimal time delay in inter-layer imaging and photostimulation, in both the mouse cortex and hippocampus in vivo. Combined with behavioral experiments, CLAOP provides all-optical evidence linking behavioral responses to neuronal connectivity in the primary visual cortex (V1) of live mice. Furthermore, we demonstrate that CLAOP can perturb the activity response of inter-layer cortical neurons to sensory stimuli according to their functional signatures. Overall, CLAOP provides an all-optical approach for mapping inter-layer connectivity at the single-neuron level and for modifying neuronal responses in behaving animals.

## Introduction

Revealing the connectome of the brain is crucial for understanding its functions and dysfunctions (Lichtman 2015; Abbott et al. 2020). Diverse approaches have been developed to dissect the anatomical or functional connectivity of individual neurons, such as single-neuron electrophysiology (Ko et al. 2011; Yoshimura et al. 2005; Song et al. 2005; Shemesh et al. 2017; Anastasiades et al. 2018) and electron microscopy-mediated tissue reconstruction (Eichler et al. 2017; Kasthuri et al. 2015; Winnubst et al. 2019; Zheng et al. 2018; Lee et al. 2016; Ding et al. 2023; Lichtman & Smith 2008). Although these approaches provide the most precise measurements for physiological or physical local connectivity, respectively, they still face challenges in high-throughput analysis with behavioral stimuli. Recently, a promising solution to address this limitation is all-optical physiology (AOP), which integrates simultaneous activity readout and manipulations within live animals (Emiliani et al. 2015; Packer et al. 2014; Marshel et al. 2019; Carrillo-Reid et al. 2016; Mardinly et al. 2018; Carrillo-Reid et al. 2019; Fan et al. 2020; Rickgauer et al. 2014; Yang et al. 2018a; Zhang et al. 2018; Daie et al. 2021; Dal Maschio et al. 2017). Combining calcium/voltage imaging with optogenetics, the AOP system can noninvasively, rapidly, and precisely acquire the local connectivity and identify the functional signature of specific neurons (Carrillo-Reid et al. 2016; Daie et al. 2021; Russell et al. 2022; Yang et al. 2018b; Spampinato et al. 2022; Carrillo-Reid et al. 2019). Moreover, AOP offers another significant advantage through its high compatibility with behavioral stimuli. When combined with behavioral approaches, the AOP strategy has the potential to generate artificially induced ensembles (Dal Maschio et al. 2017; Carrillo-Reid et al. 2016), control sensory-guided behavior (Marshel et al. 2019; Carrillo-Reid et al. 2019), and implement closed-loop control over neuronal activity patterns in awake mice (Zhang et al. 2018; Russell et al. 2022).

However, due to the limited axial switching capability of existing imaging systems, it is challenging for these all-optical approaches to dissect multi-plane functional connectivity with high temporal resolutions. The main challenge of cross-layer all-optical physiology is the loss of temporal information caused by the switching time between different axial planes. Several approaches have been developed to overcome this obstacle, such as the combination of electrically tunable lenses (Yang et al. 2018b) and ultrasound lens (Huang et al. 2019). Although these approaches have already shortened the switching time in AOP systems, more flexible and faster inter-layer methods are still required for achieving precise functional connectivity analysis, especially when employing faster activity indicators like GCaMP8 or voltage sensors. Spatiotemporal multiplexing has been utilized to enhance the temporal resolution of two-photon imaging by splitting the pulse of a femtosecond laser and introducing different optical delays to different subpulse groups (Amir et al. 2007; Cheng et al. 2011). These subpulse groups can be rearranged to different lateral and axial positions within the brain, allowing access to different neurons with only a few nanoseconds of time delay. The application of spatiotemporal multiplexing has resulted in the development of large field-of-view (FoV) two-photon imaging systems for voltage sensors (Platisa et al. 2023) and high-speed volumetric recording systems (Beaulieu et al. 2020; Demas et al. 2021). Thus, combining spatiotemporal multiplexing with AOP systems shows the potential to promote the synchronicity of recording and modulating neuronal activity, thereby facilitating high-temporal-resolution analysis of inter-layer neuronal connectivity.

In this study, we present the Cross-Layer All-Optical Physiology (CLAOP) system via integrating spatiotemporal multiplexing into a two-photon AOP system, which enables the simultaneous recording and manipulation of neural networks across brain regions with axial intervals as large as 530 μm. The temporal resolution of two-plane imaging in CLAOP can be on the scale of sub milliseconds (up to 12.5 ns temporal sampling with 1×1 pixel spatial sampling). We demonstrate the superior performance of CLAOP in interrogating inter-layer neuronal connectivity by *in vivo* imaging and manipulation of neural circuits in the cortex and hippocampus of mouse brains. We demonstrate the simultaneous recording of neurons at different layers to identify behavior-specific neurons, followed by selectively exciting these function-specific neurons while maintaining synchronous recording of neurons at multiple layers, which enables functional analysis of inter-layer connectivity. Importantly, this synchronized optical manipulation and recording of neuronal activities across long-distance layers in cortical columns *in vivo* allow us to manipulate selected neurons to achieve inter-layer modification of neural responses to behaviors. We expect that CLAOP will facilitate a broad array of novel approaches in neural connectome studies while also inspiring new applications for interacting with the brain.

## Results

### CLAOP system for simultaneous recording and manipulation of inter-layer neural networks

AOP system allows simultaneous recording and manipulation of neuronal activity (Emiliani et al. 2015; Packer et al. 2014; Marshel et al. 2019; Carrillo-Reid et al. 2016; Mardinly et al. 2018; Carrillo-Reid et al. 2019; Fan et al. 2020; Rickgauer et al. 2014; Yang et al. 2018a; Zhang et al. 2018; Daie et al. 2021; Dal Maschio et al. 2017; Oldenburg et al. 2024), enabling the dissection of activity and functional connectivity of the brain. However, local AOP has difficulties when it comes to cross-layer reading and writing. Thus, in the CLAOP system, simultaneous imaging across different planes with high temporal resolution is a major challenge. While several components, such as piezoelectric displacement stages, ETLs (Grewe et al. 2011), deformable mirrors(Shain et al. 2017), voice coils(Sofroniew et al. 2016), and ultrasonic phase-locked lenses (Kong et al. 2015, p.201), have been developed for fast axial scanning, simultaneous recording of two brain regions with significant axial separation remains difficult. To overcome this obstacle, we adopted spatiotemporal multiplexing (Amir et al. 2007; Cheng et al. 2011; Chen et al. 2016; Beaulieu et al. 2020; Demas et al. 2021; Yu et al. 2021) in our custom-built two-photon microscopes (Fig. 1A and B), which aimed to improve imaging speed between different planes. We focused one excitation beam at the aplanatic plane of the objective lens (Path 1) for optimal deep imaging quality while biasing the other beam upwards by up to 530 μm (Path 2) to cover a large axial range. The time delay between these two imaging beams is approximately 6.25 ns, thus ensuring the recording of neuronal activity with high temporal fidelity. The stimulation beam is parfocal with the imaging beam, which focuses on the deeper plane. We adopt the spiral scan strategy to photostimulate neurons expressing optogenetic opsins.

**Figure 1.**
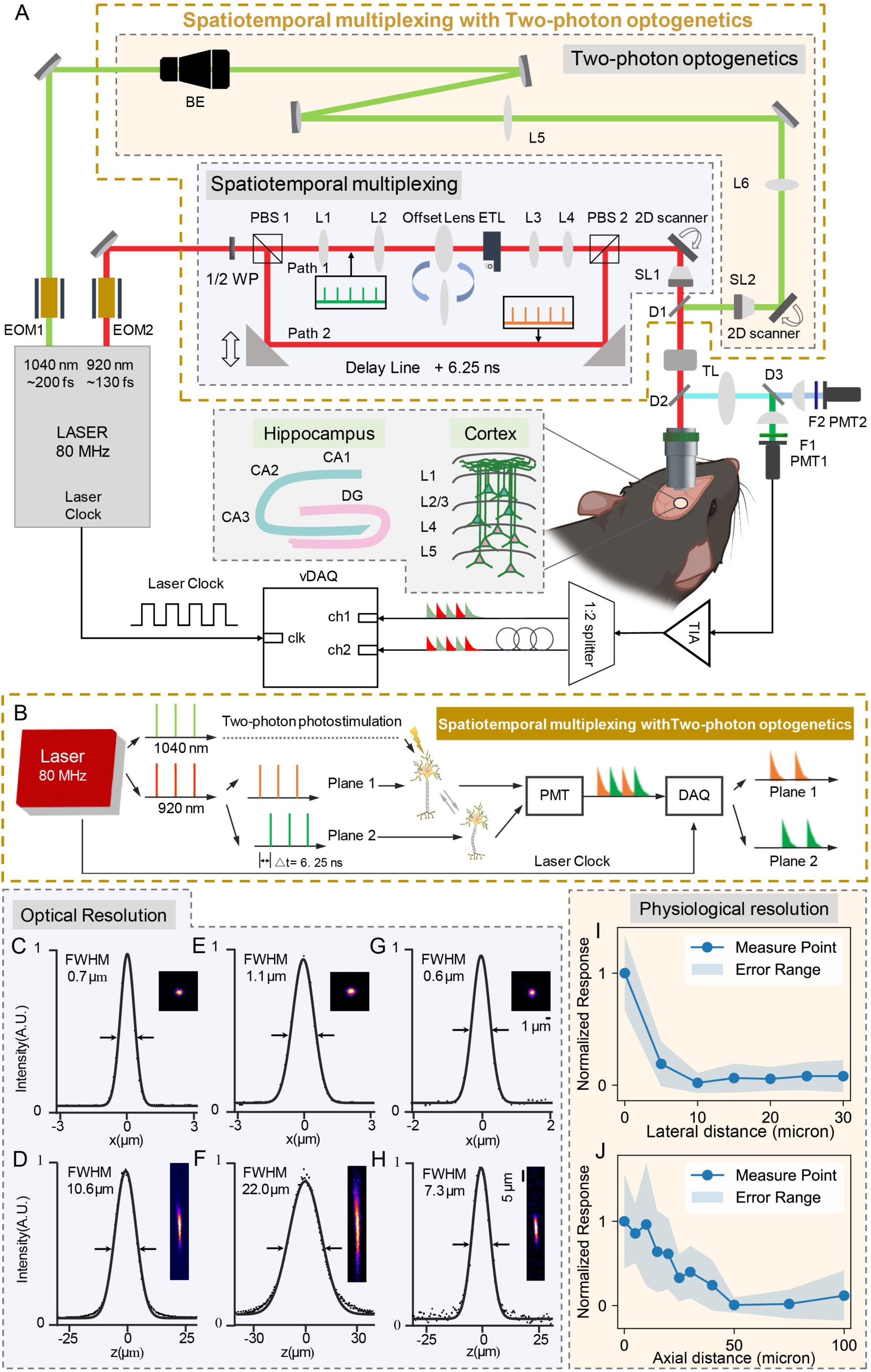
System schematics of the cross-layer all-optical physiology (CLAOP) system. **(A)** System diagram of the simultaneous multiphoton recording and manipulating system. EOM: Electro optic modulator, 1/2 WP: Halfwave plate, PBS: Polarization beam splitter, BE: Beam expander, L: Lens, SL: Scan lens, TL: Tube lens, D: Dichromatic mirror. Scale bar: 80 μm. The axial distance between the upper and lower imaging planes is adjusted by the electronic tunable lens(ETL). Optogenetic foci are located at the lower plane. **(B)** Signal flow diagram of the CLAOP system. **(C and D)** Cross-sectional plots of the focal profile in the lateral and axial direction of the lower imaging plane, respectively. FWHM full width at half maximum. **(E and F)** Cross-sectional plots of the focal profile in the lateral and axial direction of the upper imaging plane, respectively. **(G and H)** Cross-sectional plots of the focal profile in the lateral and axial direction of the photostimulation beam, respectively. **(I)** Normalized stimulation response to the photostimulation versus lateral distance between stimuli point and measured cell. ΔF/F is normalized to the averaged response of on-target stimuli. **(J)** Normalized stimulation response to the photostimulation versus axial distance between stimuli point and measured cell. ΔF/F is normalized to the averaged response of on-target stimuli. Data are represented as mean ± standard deviation of the mean.

To record neural activities at multiple cortical layers, we further inserted an electrically tunable lens (EL-16-40-TC, Optotune) into Path 2 to rapidly switch imaging planes. Moreover, by combining offset lenses of 75 mm focal length with objective lenses (CFI75 LWD 16X W, Nikon), axial imaging ranges as large as 530 μm can be achieved. We quantified the two-photon spatial resolution using 0.5 μm fluorescence beads. The lateral full-width-at-half-maximums (FWHMs) are 0.7 ± 0.1 µm (mean ± s.d.) and 1.1 ± 0.1 µm at the lower plane (Path1) and the upper plane (Path2), respectively, while the axial FWHMs are 10.6 ± 0.1 µm and 22.0 ± 0.5 µm at the lower plane (Path1) and the upper plane (Path2, measured at +480 µm depth with CFI75 LWD 16X W), respectively (Fig. 1C to F). Both imaging beams are underfilled to raise penetration depth and reducing the aberration of ETL. In comparison, the lateral and axial FWHMs of the stimulation path are 0.6 ± 0.1 µm and 7.3 ± 0.4 µm, respectively (Fig. 1 G and H), as the stimulation path is of higher numerical aperture (NA) to improve the spatial localization and photostimulation efficiency of two-photon optogenetics. As the photostimulation focus moves away from the target neuron laterally or axially, the response of the target neuron decreases(Fig. 1I and J). Overall, CLAOP demonstrates a unique and reliable capability for interrogating the functional connectivity between neurons located at different depths *in vivo*.

### High-speed cross-plane imaging of neuronal activity *in vivo* via CLAOP

To reveal activity patterns and coactivities across different planes in the brain, it is crucial to simultaneously image two axially separated layers of neurons at high temporal resolutions. First, we performed *in vivo* dual-plane imaging of the mouse V1 cortex expressing GCaMP6s at two different depths (100 μm and 530 μm) with a 540 × 540 μm^2^ FoV at 60 Hz in anesthetized mice (Fig. 2A and B) (See Table S1 for a summary of imaging conditions for all experiments). This enabled the detection of a substantial population of active neurons from both planes, facilitating further analysis of population activity patterns and coactivities among neurons within and across planes. Neurons exhibiting high cross-correlations of calcium activities were identified, suggesting potential functional connectivity (Fig. 2C to E). Next, we further demonstrated the dual-plane recording of neuronal activities from the deep and superficial layers of hippocampal CA1 expressing GCaMP8m(Zhang et al. 2023), with the overlying cortex tissue removed. Despite these two layers being separated by only 50 μm, we could separate the activity traces of neurons from both layers with high quality (Fig. 2F to H) and identify possible functional connections between neurons in CA1 via activity correlation (Fig. 2I and J).

**Fig. 2.**
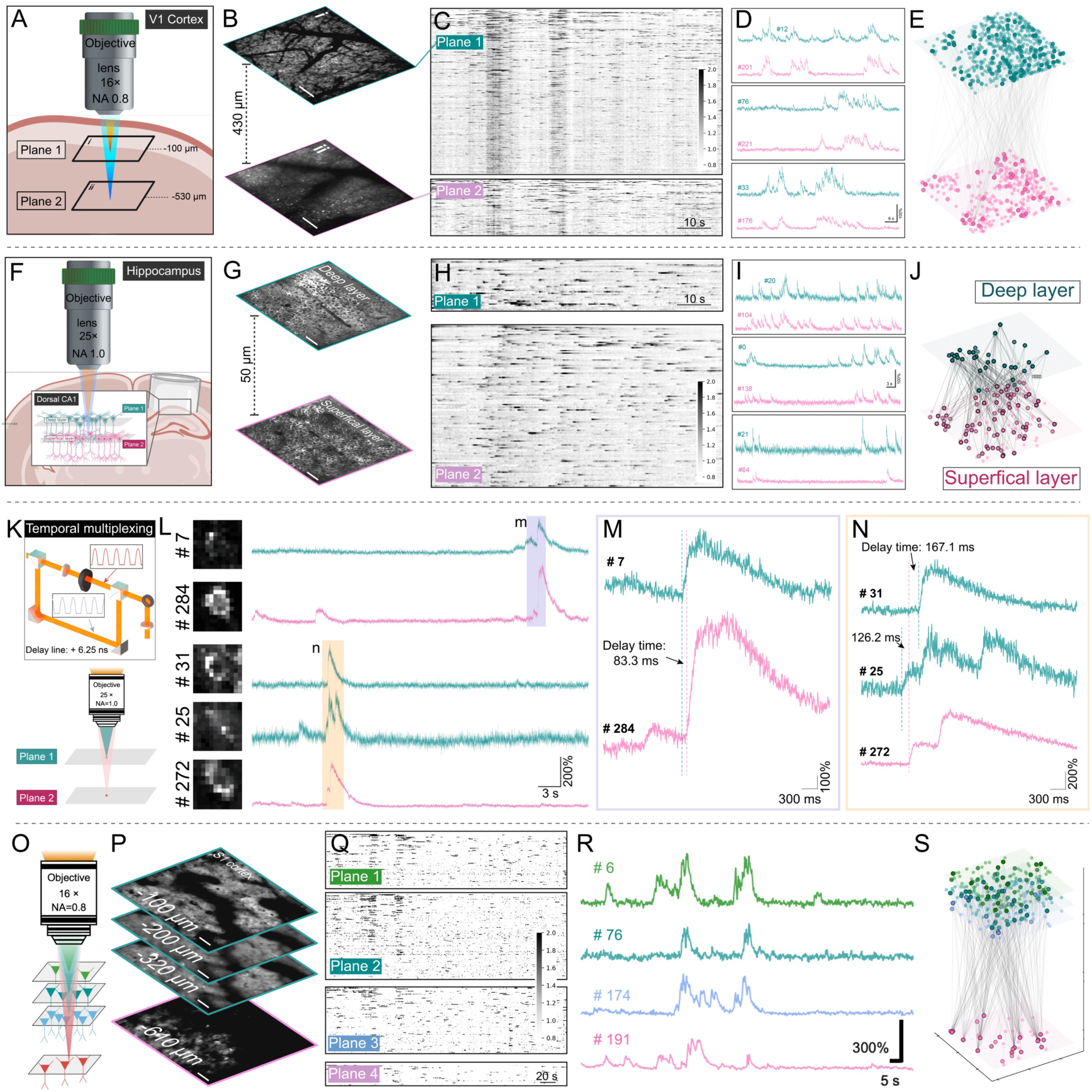
Simultaneous dual-plane recording of mouse neuronal activity. (**A**) Diagram of dual-plane imaging in the mouse V1 cortex. (**B**) Representative images of two planes simultaneously acquired in the mouse V1 area. Scale bar: 90 μm. (**C**) Neural activities of recorded neurons in both regions. (**D**) Activities of selected neurons from the dual plane. (**E**) 3D demonstration of all recorded neurons from dual-plane recording. Lines connecting neurons indicate a high correlation in activity (Corr>0.55) (**F**) Diagram of dual-plane imaging in mouse hippocampal CA1. (**G**) Representative images of two planes simultaneously acquired in mouse hippocampal CA1. Scale bar: 35 μm. (**H**) Neural activities of recorded neurons in both regions. (**I**) Activities of selected neurons from the dual plane. (**J**) 3D demonstration of all recorded neurons from dual-plane recording. Lines connecting neurons indicate a high correlation in activity (Corr>0.30) (**K**) Diagram of high frame rate (>396 Hz) dual plane imaging in mouse cortex (Axial interval: 50 μm). (**L**) Activities of selected neurons with similar activity traces from the dual plane. (**M**) Zoomed-in trace of neurons highlighted purple in (L). (**N**) Zoomed-in trace of neurons highlighted yellow in (L). (**O**) Diagram of four-plane imaging in the mouse S1 cortex. (**P**) Representative images of the four planes acquired in the mouse S1 cortex. Scale bar: 40 μm. (**Q**) Neural activities of recorded neurons in four layers. (**R**) Activities of selected neurons from four layers. (**S**) 3D demonstration of all recorded neurons from four-plane recording. Lines connecting neurons indicate a high correlation in activity (Corr>0.55).

The high temporal precision in CLAOP is ensured by a delay of fewer than 10 ns between two imaging planes via utilizing spatiotemporal multiplexing. We used the CLAOP system to perform high-speed dual-plane cortex calcium imaging (396.1 Hz) and identify the sequence of firing among highly correlated neurons from two neuronal layers (Fig. 2K to N). Using our system, we could identify the short delay times between the onset of calcium events in neurons from different planes (Fig. 2M and N). We also performed high-speed recording (60 Hz) of local GABA signals(Kolb, Ilya et al. 2022) from two neuronal layers, further demonstrating the capability of our system to record diverse imaging modalities with a high frame rate (Fig. S4).

Additionally, with the employment of spatiotemporal multiplexing, the CLAOP system is also compatible with the ETL method. By combining a switchable offset lens before the ETL, our system was further improved to achieve interlayer imaging between multiple planes. Specifically, we performed multi-plane imaging in mouse S1 cortex spanning over 500 μm, resulting in near-volumetric imaging and analysis of coactivities in a large number of neurons in a 3D volume (Fig. 2O to S). Our design led to a frame rate of 30 Hz in the lower plane and a volume rate of 10 Hz in the upper volume. These results demonstrated the versatility of the CLAOP system for studying a variety of brain regions and for different scenarios where high-speed or multi-plane sampling is required.

### Identification of single-neuron functional connectivity between different neuronal layers via CLAOP

The combination of two-photon imaging and optogenetics is a powerful strategy for dissecting functional connectivity patterns among neurons (Packer et al. 2014; Marshel et al. 2019; Rickgauer et al. 2014; Yang et al. 2018a; Zhang et al. 2018). However, most systems are still limited when measuring neuronal connectivity across axially spaced layers, which is a key question in neural circuit studies (Adesnik & Naka 2018; Nagayama et al. 2014). CLAOP system allows simultaneous dual-plane imaging and single-cell optogenetics, thus it is competent in dissecting across-layer functional connectivity. To demonstrate this potential, we first performed CLAOP on the mouse hippocampus CA3 region. Calcium imaging of neurons at two layers was recorded at 60 Hz, while optogenetic stimulation of selected neurons expressing ChRmine(Marshel et al. 2019) was performed in the lower plane (Fig. 3A and B)(See Table S1 for a summary of imaging and photostimulation conditions for all experiments). An axial distance of 50 μm was chosen between the two planes, since the off-target effect is minimal at this level (see Fig.1J and discussion). Optogenetic activation of neurons in the deeper position can elicit activity of other non-targeted neurons, consistent with studies showing the existence of recurrent connections between CA3 neurons(Miles & Wong 1986; Guzman et al. 2016) (Fig. 3C and D). We also used the CLAOP system to probe interlayer functional connectivity in the mouse S1 cortex (Fig. S5) and CA1 (Fig. S6). Notably, correlation structures generated only from recorded spontaneous activity between neurons generally could not persist across different time points. However, optogenetic stimulation-mediated coactivation patterns remained stable (Fig. S7), which indicates that CLAOP-mediated analysis of functional connectivity is more reliable, compared to analysis based on calcium recording only(Cohen & Kohn 2011).

**Fig. 3.**
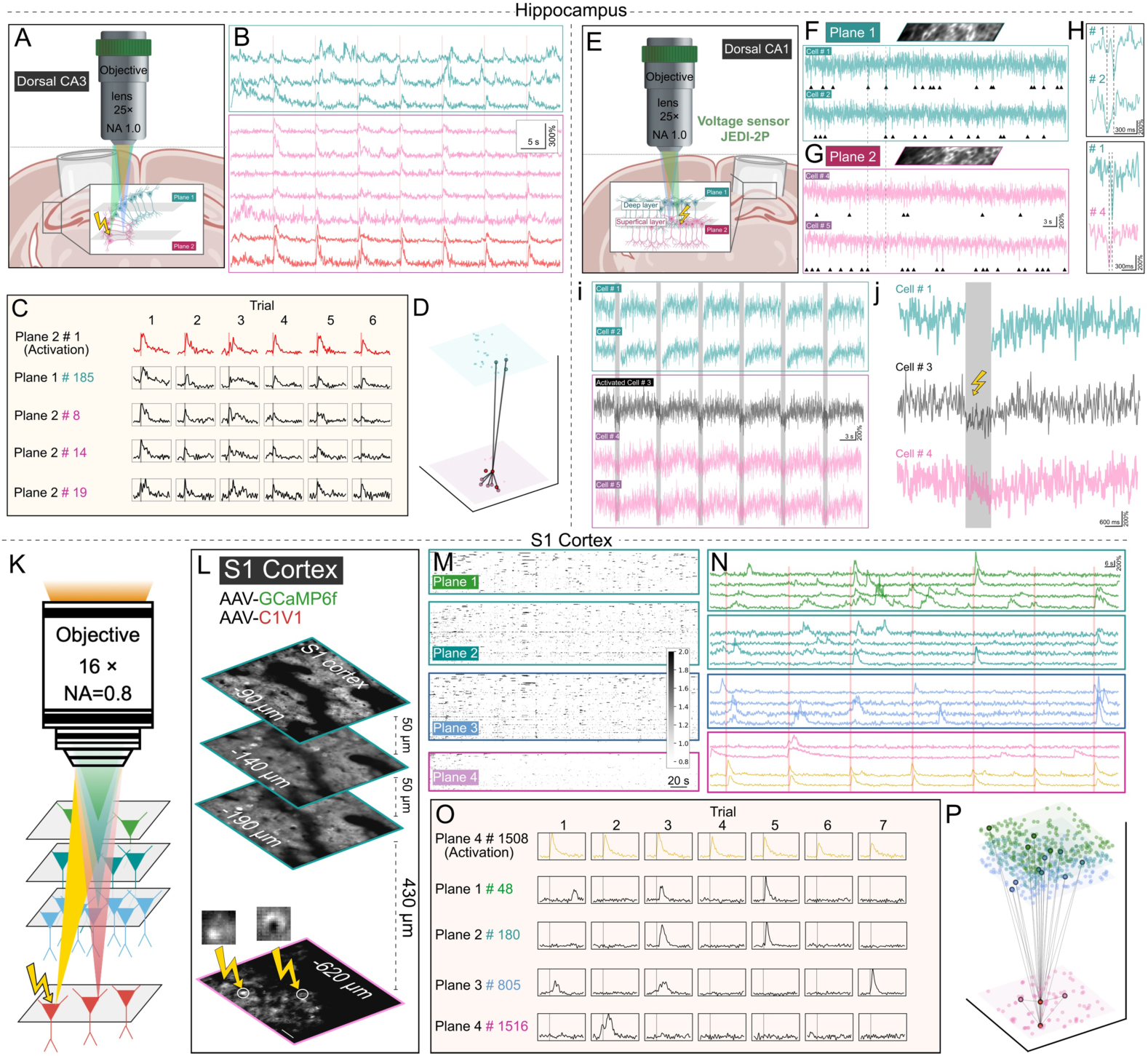
Dual-plane and 3D recording combining optogenetic manipulation. (**A**) Diagram of dual-plane recording with optogenetic manipulation in mouse hippocampal CA3 (Axial interval: 50 μm). (**B**) Activities of selected neurons from the dual plane. (cyan: upper plane, pink: lower plane, red: target cell) (**C**) Responses of the target neuron and other neurons to two-photon photostimulation. (**D**) 3D demonstration of all recorded neurons from dual-plane recording with optogenetic manipulation. Lines connecting neurons indicate a high correlation between other neurons shown in (B) and the target neuron. (**E**) Diagram of dual-plane recording with optogenetic manipulation in mouse hippocampal CA1 expressing the voltage indicator (Axial interval: 50 μm). (**F**) Representative voltage signal traces of neurons in the deep layer. Fov: 160×30 μm. Arrows indicate putative spikes. (**G**) Representative voltage signal traces of neurons in the superficial layer. Fov: 160×30 μm. Arrows indicate putative spikes. (**H**) Zoomed-in trace of neurons in (F) and (G). (**I**) Voltage traces of neurons in two layers with optogenetic activation. (J) Zoomed-in traces of neurons in (I). (**K**) Diagram of four-plane recording with optogenetic manipulation in the mouse S1 cortex. (**L**) Representative images of four planes acquired in the mouse S1 cortex with optogenetic manipulation of two target cells indicated with white circles. Scale bar: 40 μm. (**M**) Neural activities of recorded neurons in four planes. (**N**) Activities of selected neurons from four planes. (**O**) Responses of the target neurons and other neurons to two-photon photostimulation. (**P**) 3D demonstration of all recorded neurons from four-plane recording. Lines connecting neurons indicate a high correlation between other neurons shown in (N) and the target neurons.

Taking advantage of our system in inter-layer measurement at high temporal resolution, we tested the potential of CLAOP in detecting the voltage indicator JEDI-2P(Liu et al. 2022) and performed optogenetics with C1V1(Yizhar et al. 2011) in CA1 (Fig. 3E). We first demonstrated that the negative-going transients of JEDI-2P signals represent significant firing events by comparing to the calcium events reflected by jRGECO signals (fig. S9). Spiking events of neurons from two layers can be precisely visualized and aligned for detailed analysis of cofiring (Fig. 3F to H). Photostimulation-induced depolarizing events on targeted neurons and spiking events on other neurons could also be detected (Fig. 3I and J), showing the advantage of CLAOP in interrogating neural connectivity via sensitive indicators. To dissect neuronal connectivity from multiple layers, we next employed ETL in our system to perform 3D imaging, while simultaneously recording and manipulating on a separate plane (Fig. 3K). The imaging is of 30 Hz frame rate at the deeper plane and of 10 Hz volume rate at the upper volume. In the mouse S1 cortex expressing GCaMP6f and C1V1 (Fig. 3L to N), we identified non-stimulated neurons that responded to the photostimulation event, implying possible functional connectivity between the targeted neurons (Fig. 3O and P). This strategy effectively extends the pool of recorded neurons and is suitable for multi-layer connectivity dissection when utilizing fast and sensitive activity indicators.

### Analysis of functional connectivity between behavior-responsive neurons in mouse V1 cortex

In light of the promising potential of CLAOP for mapping inter-layer functional connectivity, we next combined connectivity mapping with behavioral analysis. Previous studies have utilized neuronal activity imaging and various connectivity mapping techniques, such as electrophysiology(Ko et al. 2011; Cossell et al. 2015), electron microscopy(Ding et al. 2023; Lee et al. 2016), and rabies tracing(Wertz et al. 2015; Rossi et al. 2020), to investigate whether neurons with similar functional responses are more likely to connect in the visual cortex. In this study, we employed the CLAOP system to address this question specifically in the mouse V1 cortex.

Here, we expressed GCaMP8m and red-shifted ChRmine (rsChRmine) (Kishi et al. 2022) to enable simultaneous recording and manipulation of neuronal activity *in vivo*. rsChRmine was used in this experiment to minimize the artefact activation of opsin by the imaging laser (920 nm) . Visual stimuli of drift grating were first presented to the mouse while the system recorded neuronal activities at two layers separated by 50 μm (Fig. 4A to D). Next, we photostimulated selected neurons and recorded the responses of all neurons from both layers (Fig. 4E to G). We observed a positive correlation between the coupling strength of neuronal pairs and their activity correlation during visual stimulation (Fig. 4H and I). This finding is consistent with previous studies demonstrating that functional activity correlation or similar stimulus selectivity positively correlates with the likelihood of neuronal connectivity at various levels (Wertz et al. 2015; Rossi et al. 2020). A similar result was also obtained by CLAOP when mice were given movies as visual stimuli (Fig. S8). Thus, our method allows a broader AOP analysis of both axial and lateral connectivity.

**Fig. 4.**
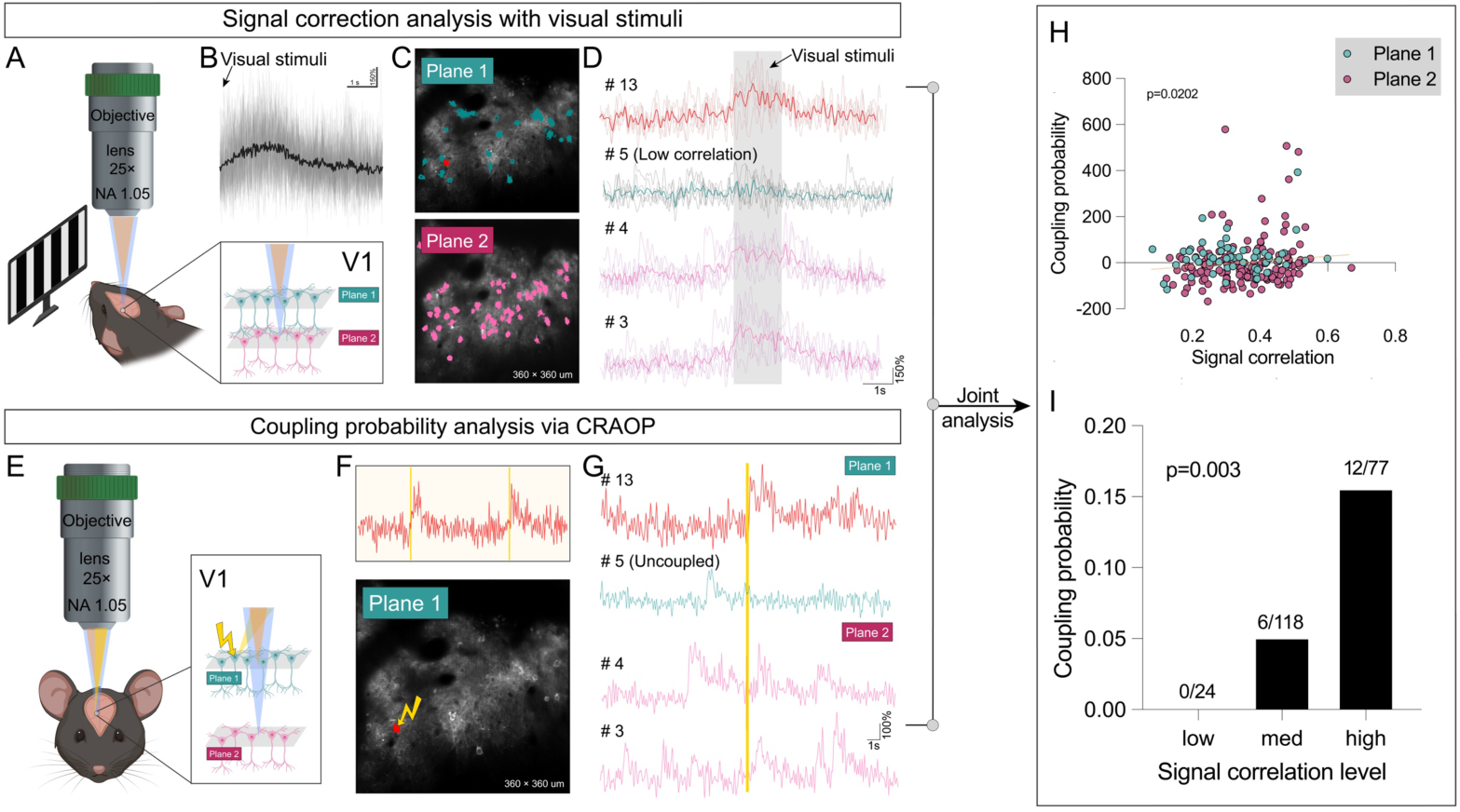
CLAOP analysis of functional connectivity with stimulus response correlation in V1. (**A**) Diagram of dual-plane recording in V1 with visual stimuli. (**B**) Averaged activity trace of recorded neurons in response to the visual stimuli. (**C**) Example of spatial masks of recorded neurons in one trial (cyan: upper plane cells, pink: lower plane cells, red: cell to be photostimulated). (**D**) Example of activity traces of selected neurons (#13 is the cell to be stimulated; #5 has a low signal correlation with it; #3 and #4 have a high signal correlation with it). (**E**) Diagram of dual-plane recording and targeted photostimulation in V1. (**F**) Example activity trace of the stimulated cell. (**G**) Example of activity traces of selected neurons. (#13 is the stimulated cell; #5 has a low coupling strength with it; #3 and #4 have a high coupling strength with it). (**H**) Linear regression plot. The coupling strength of neuron pairs was fitted as a function of the signal correlation between the pair. Regression coefficient = 143.92. P=0.0202. (**I**) A significant increase in coupling probability with increasing signal correlation to visual stimuli (P=0.003, Cochran–Armitage test for trend).

### CLAOP-mediated artificial modification of neuronal response in mouse V1 cortex

Building upon its efficacy in mapping neuronal connectivity, our system extends its capabilities to manipulate neuronal activity, thereby enabling targeted perturbations within the neural network. To demonstrate its effectiveness, our investigation focused on the V1 cortex. We aimed to investigate whether perturbing visual-responsive neurons alters the function of other non-targeted neurons from different layers. Mice were given drifting grating visual stimuli while dual-plane calcium imaging was performed in the V1 area, covering a superficial layer of neurons and a deeper layer of neurons (Fig. 5A). Visually responsive cells were identified in both Plane 1 (130 μm below the brain surface) (Fig. 5B to D) and Plane 2 (330 μm below the brain surface) (Fig. 5E to G). Subsequently, selected visually responsive neurons in the deeper layer were photostimulated during visual stimuli presentation. Specifically, the neurons were activated when nonpreferred orientation stimuli were presented, aiming to perturb their normal visual responsiveness (Fig. 5H). Non-targeted neurons in both Plane 1 and Plane 2 exhibited a change in orientation selectivity during these trials (Fig. 5I to N). The targeted neurons indeed exhibited increased activity after photostimulation, resulting in a change in selectivity (Fig. 5K). Additionally, some non-targeted neurons displayed altered visual responses at the photostimulation time points (Fig. 5M). Notably, the impact of the perturbation on some non-targeted neurons caused significant disparity in the time points of photostimulation and the subsequent substantial change in their activity (Fig. 5L). This may be explained by the involvement of intricate local circuits and computational processes in visual information processing within the V1 cortex (Cossell et al. 2015). These results showcase the potential of the CLAOP system to manipulate and record neuronal activity across distinct axially separated layers. The integration of connectivity mapping, single-cell manipulation, and recording using the CLAOP system, along with behavioral analysis, can provide deeper insights into the local circuit architecture and cortical computation mechanisms.

**Fig. 5.**
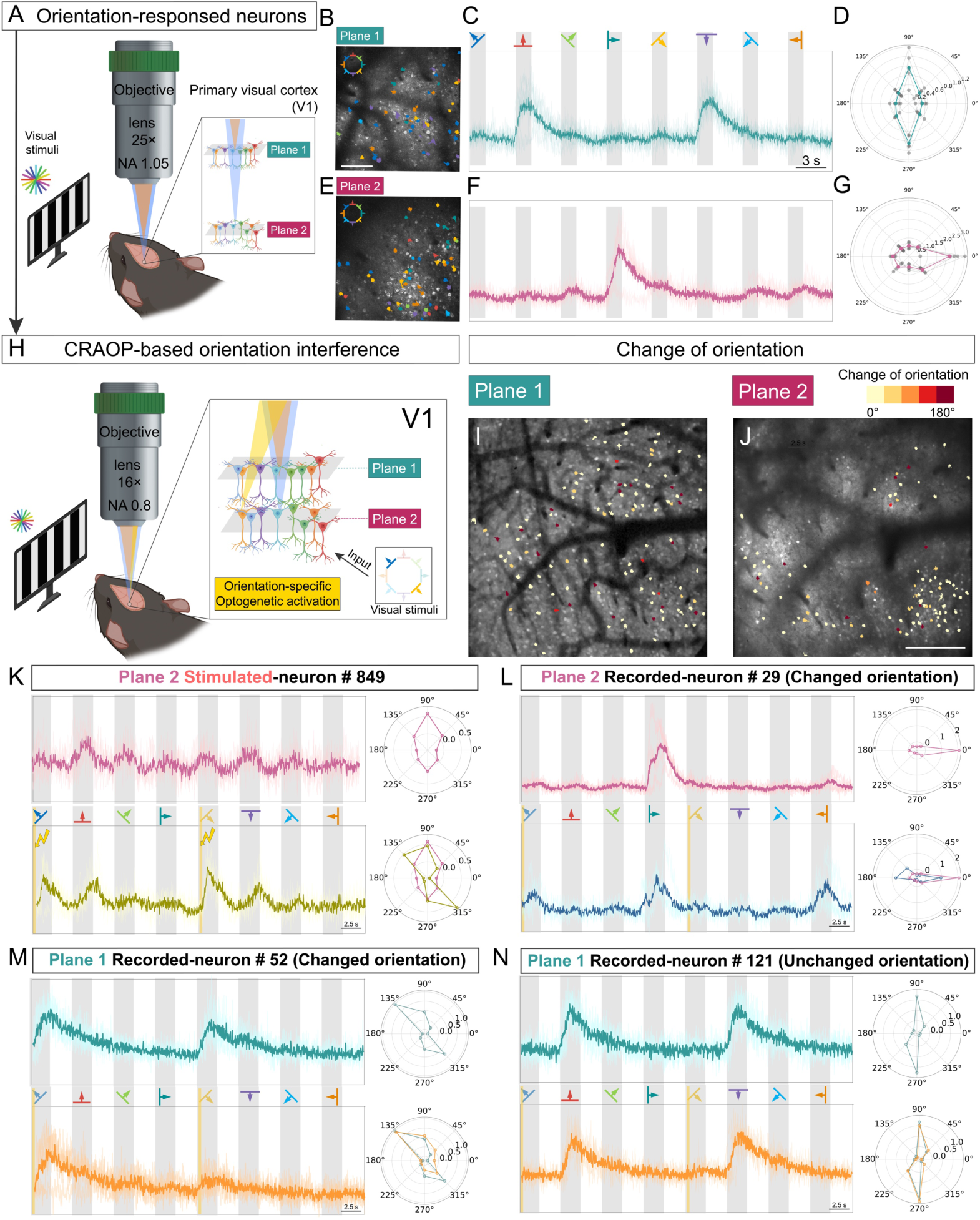
Simultaneous dual-plane recording and optogenetic perturbation of mouse V1 neuronal activity during visual stimuli presentation. (**A**) Diagram of dual-plane recording under drifting grating visual stimuli. (**B**) Orientation selectivity map of plane 1 (140 μm depth). Scale bar: 100 μm. (**C** and **D**) Activity trace and corresponding polar plot of representative visual response neurons in plane 1. (**E**) Orientation selectivity map of plane 2 (240 μm depth). (**F** and **G**) Activity trace and corresponding polar plot of representative visual response neurons in plane 2. (**H**) Diagram of dual-plane recording with photostimulation under drifting grating visual stimuli. (**I** and **J**) Orientation change pattern before and after photostimulation (130 μm and 330 μm depth). Scale bar: 150 μm. (**K**) Activity trace and corresponding polar plot of the target neuron in plane 2. (**L**) Activity trace and corresponding polar plot of a nontarget neuron in plane 2. (**M** and **N**) Activity traces and corresponding polar plots of nontarget neurons in plane 1.

## Discussion

An effective strategy to achieve functional connectivity analysis relies on simultaneous readout and manipulation of neuronal activity during the presentation of behavioral stimuli. While electrophysiology or optogenetics combined with calcium imaging can precisely analyze local circuits(Marshel et al. 2019; Carrillo-Reid et al. 2019; Fan et al. 2020; Packer et al. 2014; Zhang et al. 2018), mapping inter-layer connectivity remains necessary. In this work, we demonstrated a cross-layer all-optical physiological approach with a high temporal resolution, showing the unique advantages in the dissection of inter-layer functional connectivity. Overall, our system represents a significant improvement in interrogating inter-layer connectivity with high spatial and temporal precision *in vivo*.

### Comparison of CLAOP with other all-optical physiology systems

Compared to previously reported three-dimensional AOP systems(Mardinly et al. 2018; Dal Maschio et al. 2017; Huang et al. 2019; Yang et al. 2018b), our system offers a significant advantage in terms of temporal resolution for cross-plane recording and optogenetic manipulation. In our system, the adoption of spatiotemporal multiplexing(Amir et al. 2007; Cheng et al. 2011; Chen et al. 2016; Beaulieu et al. 2020; Demas et al. 2021; Yu et al. 2021) ensures simultaneous two-photon imaging of multiple focal planes within live mice, minimizing time delays among different planes (6.25 ns). This enables a more precise analysis of activity correlation between upstream and downstream neurons following optogenetic stimulation. Moreover, our system also supports simultaneous high-frame-rate imaging at different planes, making it appropriate for fast and sensitive activity indicators such as GCaMP8(Zhang et al. 2023) and voltage sensors(Liu et al. 2022; Peterka et al. 2011; Vogt 2015). In addition, the spatiotemporal multiplexing technique used to expand the imaging voxel speed limits the number of multiplexed beams by the fluorescence lifetime. Our system also has the potential to benefit studies of continuous spatial dynamics of biological factors in the brain, which also require high-temporal-resolution 3D imaging, such as the motion of glia, blood flow, and dynamic distribution of molecules within cells. Nonetheless, our results show that CLAOP is compatible with other fast axial scanning devices, such as ETL, to further promote the ability for multi-plane all-optical physiological measurement (see Figs. 2 and 3).

### The limitations of the CLAOP system

To achieve precise population-level connectivity analysis and closed-loop manipulation of behavioral output, it is possible to improve our CLAOP system by integrating other techniques, such as anatomical analysis (The MICrONS Consortium et al. 2021), large-field-of-view imaging (Sofroniew et al. 2016; Kim et al. 2016), and more efficient neuronal manipulation like 3D-SHOT (Pégard et al. 2017). Specifically, by implementing holographic optogenetics in our system, it will be possible to test how perturbations on a neuronal ensemble, but not single neuron, affect the functional connectivity across neuronal layers and even behavioral outputs (Carrillo-Reid et al. 2019; Marshel et al. 2019; Zhang et al. 2018). Currently, our imaging system emphasizes the fast all-optical analysis of neurons across different layers rather than distant lateral regions in mouse brain imaging. Future enhancements will involve integrating a meso-objective lens with dual scan engines, thereby expanding the capacity of the CLAOP system for cross-region interrogation of neuronal connectivity (Yu et al. 2021). Additionally, temporal focusing has been utilized in optogenetic manipulation to enhance the axial physiological resolution of photostimulation (Faini et al. 2023; Pégard et al. 2017). Therefore, integrating CLAOP with temporal focusing, which offers more confined axial excitation, will provide more reliable connectivity mapping in densely arranged brain regions, such as hippocampus CA1 and DG.

## Materials and Methods

### Optical setup of the CLAOP system

The detailed system setup is demonstrated in Fig. 1. The system is composed of a pulsed femtosecond laser (80 MHz, Chameleon Discovery, Coherent) which has two outputs and a custom-built two-photon laser scanning module. The imaging laser which has a built-in dispersion compensation module is tuned to 920 nm for calcium imaging. The stimuli laser is fixed to 1040 nm to perform two-photon optogenetics stimulation. The imaging laser power is controlled with a Pockels cell (Pockels cell: 350-80-LA, Controller: 302RM, Conoptics, Inc.). After the Pockels cell, a half-wave plate (AHWP05M-980, Thorlabs) mounted on an electric rotation stage (PRM1/MZ8, Thorlabs) and a polarization beam splitter (PBS203, Thorlabs) are introduced into the optical path to realize adjustment of energy distribution between two imaging planes. The lower plane imaging path is ∼1.8 m shorter than the upper plane imaging path which leads to ∼6.25 ns delay between two beamlets. The upper plane imaging path goes through a 2X beam expander composed of two achromats (AC254-75-B and AC254-150-B, Thorlabs) to expand beam width. An offset achromat (AC254-75-B\AC254-250-B, Thorlabs), an ETL (EL-16-40-TC, Optotune), and two other achromats (AC254-200-B and AC254-100-B, Thorlabs) are placed at different intervals to obtain designed focal power bias. The lower plane imaging beam and upper plane imaging beam are combined with another polarization beam splitter (CCM1-PBS253/M, Thorlabs). A 6-mm galvanometric scan mirror (8315K, Cambridge Technology) and an 8 kHz Resonant mirror (CRS 8 kHz, Cambridge Technology) lead to a 60Hz imaging framerate at 256×256 pixel number. The combined beam is then imaged to the objective (16X CFI75 LWD, NA 0.80, Nikon) pupil by the combination of a scan lens (SL50-2P2, Thorlabs) and a tube lens (TL200-CLS2, Thorlabs). The objective is mounted on a piezo stage (P-725.4CD, Physik Instrumente) to perform long-range Z-scanning. A three-dimensional (3D) high-precision motor stage (Stage: M-VP-25XA-XYZL, Controller: ESP301, Newport) is used to move the sample relative to the objective.

In the 1040 nm photostimulation path, another Pockels cell (Pockels cell: 350-80-LA, Controller: 302RM, Conoptics, Inc.) is used to modulate the photostimulation power. An X-Y 2-axis galvanometric scan mirror (8315K, Cambridge Technology) steers the beam to perform a serial spiral scan of neurons. After a scan lens (SL50-2P2, Thorlabs), the stimuli beam is combined with the imaging beam by a dichroic mirror (DMSP1000R, Thorlabs).

The fluorescence signal is separated from the main excitation path by a dichroic mirror (DMLP805R, Thorlabs). A collection lens pair (#49-284, Edmund; KPA034-C, Newport) collects as many possible photons to enhance system efficiency. Another dichroic mirror (FF560-Di01-35.5×50.0, Sermock) split the emission light into two channels in which red (FF01-632/148-29-D, Sermock) and green (FF03-525/50-30-D, Sermock) fluorescence emission filters are located. Photomultiplier tubes (H10770PA-40, Hamamatsu) are used to detect emission light. The detection arm design is mainly based on MIMMS (Janelia Farm Research Campus). The system is controlled by scanimage (vidiro) software(Pologruto et al. 2003).

We replaced the offset achromat (AC254-75-B, Thorlabs) with another achromat (AC254-250-B, Thorlabs) in the experiment with the axial interval between two planes less than 300 microns.

For multi plane imaging with ETL demonstrated in Fig.2 O to S and Fig.3 K to P (Table S1), we collect 256 x 512 pixels images which leads to a frame rate of 30 Hz in the lower plane and a volume rate of 10 Hz in the upper volume. And 50% pixels were kept to avoid override effect of ETL.

### Signal demultiplexing of the CLAOP system

The wiring diagram of the data acquisition system is demonstrated in Fig. 1. A high-speed trans-impedance amplifier (DHPCA-100, Femto) amplifies current signals from PMT and sends voltage signals for subsequent demultiplexing. After amplification and filtering, fluorescence signals are split into two channels by a power splitter (ZAPD-30-S+, MiniCircuits)(Xiao & Mertz 2019). A longer BNC cable (∼1.3 m) delays one of the channels relative to another by 6.25 ns. We then synchronize the data sampling clock of the data acquisition board (vDAQ data acquisition hardware, vidiro) to the clock pulse train generated by femtosecond laser. To sample each channel at the peak of the signals, the laser clock phase is also adjusted by altering the length of the BNC cable according to the intensity of the image. Thus, the peak of fluorescence signal pulse emitting from the lower plane is captured at channel 1. Accordingly, the peak of the fluorescence signal pulse emitting from the upper plane is captured at channel 2. The bandwidth of the transimpedance amplifier can be adjusted to strike a balance between signal-to-noise ratio and signal aliasing (Fig. S1 and 2). The remaining signal mixing between two channels can be solved by linear demixing(Chen et al. 2016) (Fig.S3).

### Spatial resolution calibration of the CLAOP system

The experimental resolution of the microscope is evaluated by measuring the excitation point spread function (PSFex) as the full width at half-maximum (FWHM) of the imaging intensity profile of 0.51 µm green fluorescent polymer microspheres (Fluoro-Max, Dia: 0.51 µm, Thermo Fisher Scientific) buried in 2% agarose gel. Cross sections of green fluorescent polymer microspheres are also illustrated in Fig. 1.

### Optogenetics calibration of CLAOP system

Since the opening of the opsin ion channel is a cumulative effect, and the axonal dendrites adjacent to the soma may also be located in the path of the excitation light, even the soma that is not located in the precise focal point still has a certain probability of being activated. Stray light or poor target selection can also contribute to this neighboring associated excitation. To evaluate this putative ‘off-target’ stimulation from our optogenetics manipulation, we quantify the relative response of opsin-expressed neurons with different targeted locations. Ten seconds of data pre-stimulus is used to calculate the baseline value. The maximum response of two seconds of data after the optogenetic stimulus is used to calculate the ΔF/F. The stimulus patterns are spaced away from the soma at certain intervals. The data from cells (N = 4) are finally normalized to the mean value of the excitation peaks of each cell (Fig. 1 I). Measurement results show that photostimulation in our system has high lateral resolution. Axial ‘off target’ stimulation is also measured. ΔF/F response is calculated from cells (N = 4) and normalized to the mean value of the excitation peaks (F of each cell when the photostimulation focus is at the same axial position with the imaging beam (Fig. 1 J).

### Visual stimuli setup and analysis

A liquid crystal display monitor with 1920×1080-pixel resolution is placed 15 cm away from the mouse’s eyes. The front of the screen is covered with a blue filter film to reduce visual stimuli light getting into PMT. We generated drifting sine wave gratings in different directions (0,45,90,135,180,225,270,315 degrees, 5 trials) by custom software in Psychtoolbox running on MATLAB. Each direction of drifting grating lasted 2 s and uniform gray was displayed between different directions of drifting gratings. To further reduce screen light pollution, the screen display was set to blue only. The monitor is controlled by another computer. The synchronization of visual stimuli and two-photon imaging is achieved by counting the frame trigger with a NI USB6008 DAQ card. The orientation selectivity index (OSI) is calculated as (response to preferred direction − averaged response to two orthogonal directions) / (response to preferred orientation + averaged response to two orthogonal directions)(Hagihara et al. 2015). In Fig. S8, the visual stimuli were replaced by presentations of cartoon video clips on the display monitor described above.

### Animals

All experimental procedures were approved by the Institutional Animal Care and Use Committee (IACUC) of Tsinghua University and were performed using the principles outlined in the Guide for the Care and Use of Laboratory Animals of Tsinghua University. All mice used in this study were male and imaging was performed when mice were 12-16 weeks old. All C57BL/6J mice were maintained under standard conditions by the Animal Research Center of Tsinghua University. Thy1-GCaMP6s(Dana et al. 2014) (stock number # 024275), Ai162(Daigle et al. 2018)(TIT2L-GC6s-ICL-tTA2)-D mice (stock number # 031562) and Camk2a-cre(Tsien et al. 1996) (stock number # 005359) mice were obtained from The Jackson Laboratory. Transgenic mice were genotyped through PCR using genomic DNA and Jackson Laboratories-provided primers.

### Viral constructs

The viral constructs used in the experiment were stored at -80°C until use after being subdivided into aliquots. For calcium imaging and optogenetics in CA1, a mixture of AAV_2/9_-CaMKIIα-ERCreER (OBiO), AAV_2/9_-EF1α-DIO-ChRmine-mScarlet (OBiO), and AAV_2/9_-NES-GCaMP8m (Taitool) was prepared. For calcium imaging and optogenetics in CA3, a mixture of AAV_2/9_-CaMKIIα-ERCreER (OBiO) and AAV_2/9_-EF1α-DIO-ChRmine-mScarlet (OBiO) was prepared. For calcium imaging and optogenetics in the cortex, AAV_2/9_-hSyn-C1V1(E122T/E162T)-TS-mCherry or AAV_2/9_-hSyn-NES-GCaMP8m-p2A-rsChRmine (Taitool) was used. For voltage imaging, a mixture of AAV_2/9_-CaMKIIα-Cre (OBiO), AAV_2/9_-EF1α-DIO-JEDI-2P-Kv (OBiO), and AAV_2/9_-CaMKIIα-C1V1(t/t)-ER2 (Taitool) was prepared. For the validation experiment in Fig.S9, a mixture of AAV_2/9_-CaMKIIα-Cre (OBiO), AAV_2/9_-EF1α-DIO-JEDI-2P-Kv (OBiO), and AAV_2/9_-CaMKIIα-NES-jRGECO1a was prepared. For GABA imaging, a mixture of AAV_2/9_-CaMKIIα-Cre and AAV_2/9_-EF1α-DIO-iGABAsnFR2 was prepared. The viral concentration of all viruses in the mixture was adjusted to 1-2 × 10^12^ viral genomes (v.g.)/ml by adding a buffer (PBS 1× with 5% glycerol).

### Surgery procedures

#### Hippocampus surgery

For virus injections into the hippocampus, mice were first anesthetized with Avertin (0.3g/kg body weight). Craniotomies on the skull over the right hippocampus were performed using a 0.5 mm-diameter drill, and 300 nl of the virus was injected into the dorsal CA1 (-2.0 mm A/P, -1.5 mm M/L, and −1.4 mm D/V) or CA3 (-2.7 mm A/P, - 2.0 mm M/L, and −1.9 mm D/V) using a 10 μl nanofil syringe controlled by UMP3 and Micro4 system (WPI) with a speed of 60 nl/min. After the injection, the needle remained in place for 10 min to ensure that the virus spread to the targeted area before it was slowly withdrawn. After injection, mice were returned to their home cages and were allowed to recover for at least 5 days before performing further experiments.

Imaging window implantation was performed 5 days to 14 days after the virus injection. Anesthesia was induced with 5% isoflurane before surgery and was kept with 1.5% isoflurane during the surgery. Prior to surgery, mice were administered an anti-inflammatory drug Meloxicam (8 mg/kg) subcutaneously. A 3 mm-diameter craniotomy was made using a micro-drill at the AAV injection site. The cortex tissue above the hippocampus was aspirated with a 27-gauge needle connected to a pump until the fibers of the corpus callosum became visible, exposing the alveus of the hippocampus(Sun et al. 2020; Wang et al. 2021; Ulivi et al. 2019). A stainless-steel cannula (3 mm diameter, 1.6 mm height) covered by a cover glass (3 mm diameter, 0.17 mm thickness) was inserted into the opening until the glass was in contact with the fibers. The cannula was secured in place with glue and dental cement, and a stainless-steel head-post was also fixed onto the skull using dental cement. Following surgery, mice were returned to their home cages and were monitored daily. Meloxicam (4 mg/kg/day) was given subcutaneously for 3 consecutive days after surgery to prevent inflammation. Mice were allowed to recover for at least 3 weeks before imaging experiments were performed.

#### Cortex surgery

For mouse cortex imaging experiments in Fig.1, Fig.2, and Fig.5, double transgenic Ai162/ Camk2a-cre mice were used. AAV-hSyn-C1V1(E122T/E162T)-TS-mCherry virus was injected into the targeted area. An amount of 300 nl of the virus was injected into layer 2/3 (∼250 μm deep) and layer 5 (∼500 μm deep) of the S1 (-1 mm A/P and - 3 mm M/L) or V1 (-4 mm A/P and -3 mm M/L) of the mouse cortex 3 weeks before the craniotomy surgery. The virus was front-loaded into the beveled glass pipette and injected at a rate of 50 ∼ 80 nL/min. After the injection, the needle remained in place for 10 min to ensure that the virus spread to the targeted area before it was slowly withdrawn. Craniotomy surgery was performed by drilling a 3 mm-diameter opening in the skull centered at the injection site. A circular glass coverslip was placed and sealed using a cyanoacrylate adhesive, and a stainless-steel head-post was fixed onto the skull using dental cement.

For experiments in Fig.4, C57BL/6J mice were used. First, a cranial opening was performed as described above and then AAV_2/9_-hSyn-NES-GCaMP8m-p2A-rsChRmine virus was injected into the V1 area (-4 mm A/P and -3 mm M/L). Injections were performed at a rate of 60 nl/min at 3 depths (-150 μm, -200 μm, and -250 μm below the dura surface), resulting in a total amount of 600 nl virus injected. After injection at each depth, the needle remained in place for 10 min before it was slowly withdrawn. After the virus injection, the coverslip and head-post were placed and sealed as described above.

Prior to all cortex surgeries, mice were administered Meloxicam (8 mg/kg) and Dexamethasone (1 mg/kg) subcutaneously. Meloxicam (4 mg/kg/day) was also given subcutaneously for 3 consecutive days after the surgery. Mice were used in imaging experiments at least 2 weeks after the craniotomy surgeries.

### Treatments on mice for imaging experiments

#### Sparse labeling of neurons

Tamoxifen dissolved in corn oil (5 mg/ml) was administrated to mice via intraperitoneal injection (30mg/kg body weight) 1 week before imaging. This allowed the expression of ChRmine-mScarlet in a sparse subpopulation of CA1 or CA3 neurons (for corresponding experiments in Fig.3 and Fig. S6).

#### Induction of seizure-like behaviors

For experiments on GABA signals, mice were injected with kainic acid (5mg/kg body weight, i.p.) before imaging. Seizure-like behaviors typically appeared 30 minutes after drug injection. Imaging was performed during the seizure-like behaviors with a supplement of 0.5% isoflurane for light anesthesia.

#### Anesthesia

During the majority of imaging and optogenetics experiments, mice were head-fixed and anesthetized by 1% to 1.5% isoflurane. For visual stimulation experiments, no anesthesia was given during the visual stimuli presentation. For experiments in Fig. 4 and Fig. S8, mice were given an intraperitoneal injection of chlorprothixene (5 mg/kg) before head fixation for a sedative effect.

### Data analysis

#### Calcium data analysis

For all calcium imaging data, Suite2p was used to extract spatial footprints and activity traces of putative neurons(Pachitariu et al. 2017). The results were manually inspected and refined by removing ROIs that were misclassified as neurons by the software. Fluorescence fluctuations in the surrounding neuropil of each cell were subtracted from the raw fluorescence traces (coefficient 0.7), and then ΔF/F_0_ was calculated for each ROI. F_0_ was calculated as the mean fluorescence intensity of the 10% frames with the lowest intensity. A low-pass Butterworth filter was applied to the raw extracted trace for the calcium traces displayed in the figures.

#### Voltage data analysis

For all voltage imaging data, the voltage signal was extracted after image registration. The ROIs are determined by the morphology of neurons in the averaged image. The Z-score algorithm is used to normalize the activity trace of neurons. The baseline is calculated as the median of the activity trace.

#### GABA signal data analysis

For the GABA sensor imaging data, the signal was extracted manually from selected areas in the image after image registration. The Z-score algorithm is applied to the GABA sensor signal trace. The baseline is calculated as the median of the activity trace.

#### Calculation of coupling strength and activity correlation

For data in Fig.4, the coupling strength of a neuron to a photostimulated neuron was calculated using the following method. First, the mean fluorescence value in a 0.5- second window after the end of the photostimulus was calculated as A_i_. Next, the mean fluorescence value in a 3-second window before the onset of the photostimulus was calculated as Bi. The coupling strength for this neuron to the targeted photostimulated neuron was calculated by averaging all S_i_= A_i_ – B_i_ over all such stimulation trials.

To determine which neurons are presumably coupled with the targeted neuron, we made several assumptions. We assumed if the neuron pair is not coupled, the coupling strength should be sampled from a Gaussian distribution centered around zero due to stochastic fluctuations and firings. We also assume that strong negative coupling between neurons is rare. Thus, we set a threshold for detecting coupled neurons. The threshold is the absolute value of the 5th percentile of all negative coupling strength values in all trials in a mouse. If the coupling strength exceeds this threshold, we assume it is unlikely to purely result from stochasticity, and we denote the corresponding neuron pairs as coupled.

For correlation analysis, the rolling-averaged calcium traces were used to compute Pearson’s correlation coefficient of a pair of neurons in a given time window. For representive results in Fig.2 and 3, thresholds were selected for clearly visualizing CLAOP-indentified high correlated funcational connection between neurons. These selected thresholds do not represent any biological hypothesis. Future studies should define indentified functional connection based on specific biological context.

The signal correlation of neurons in V1 was calculated for the whole time window in which visual stimuli were presented. In Fig.4, for defining low, medium, and high signal correlation levels, the values 0.2 and 0.4 were used as boundaries. The noise correlation (Fig. S7) was calculated for the time points in which no visual stimuli were presented (when mice were in the darkness).

### Statistics

Statistical tests were performed using GraphPad Prism 9 and customized R/Python scripts. The linear regression model was used for fitting the relationship between coupling strength and signal correlation. Cochran–Armitage test for trend was used to test for the significance of a linear trend in proportions. The two-tailed Wilcoxon signed-rank test was used as a paired difference test without the assumption of normally distributed samples. A p-value less than 0.05 was considered statistically significant. ∗p < 0.05; ∗∗p < 0.01; ∗∗∗p < 0.001; ∗∗∗∗p < 0.0001; n.s., nonsignificant (p > 0.05).

## Acknowledgments

We greatly thank Prof. Song-hai Shi and Dr. Yang Lin at Tsinghua University for their aid and suggestions on the experiments. We greatly thank Dr. Ilya Kolb and Jeremy Hasseman at Janelia Research Campus for sharing the plasmid sequence of iGABASnFR2. We thank all the members of the Zhong lab and Kong lab for their support.

## Funding

This work is supported by: STI2030-Major Projects 2022ZD0212000

STI2030-Major Projects 2022ZD0204900

National Natural Science Foundation of China (NSFC) 32021002

National Natural Science Foundation of China (NSFC) 61971265

“Bio-Brain+X” Advanced Imaging Instrument Development Seed Grant

## Author contributions

Conceptualization: YZ, LK, BL

Methodology: CL, YH, LK, BL

Investigation: YH, CL

Visualization: CL, YH, BL

Supervision: YZ, LK, BL

Writing—original draft: CL, YH, LK, BL

Writing—review & editing: CL, YH, YZ, LK, BL

## Data and materials availability

The data that support the findings of this study are available from the corresponding author upon reasonable request. The code used in this study is available from the corresponding author upon reasonable request.

## Figures

**Fig. S1.**
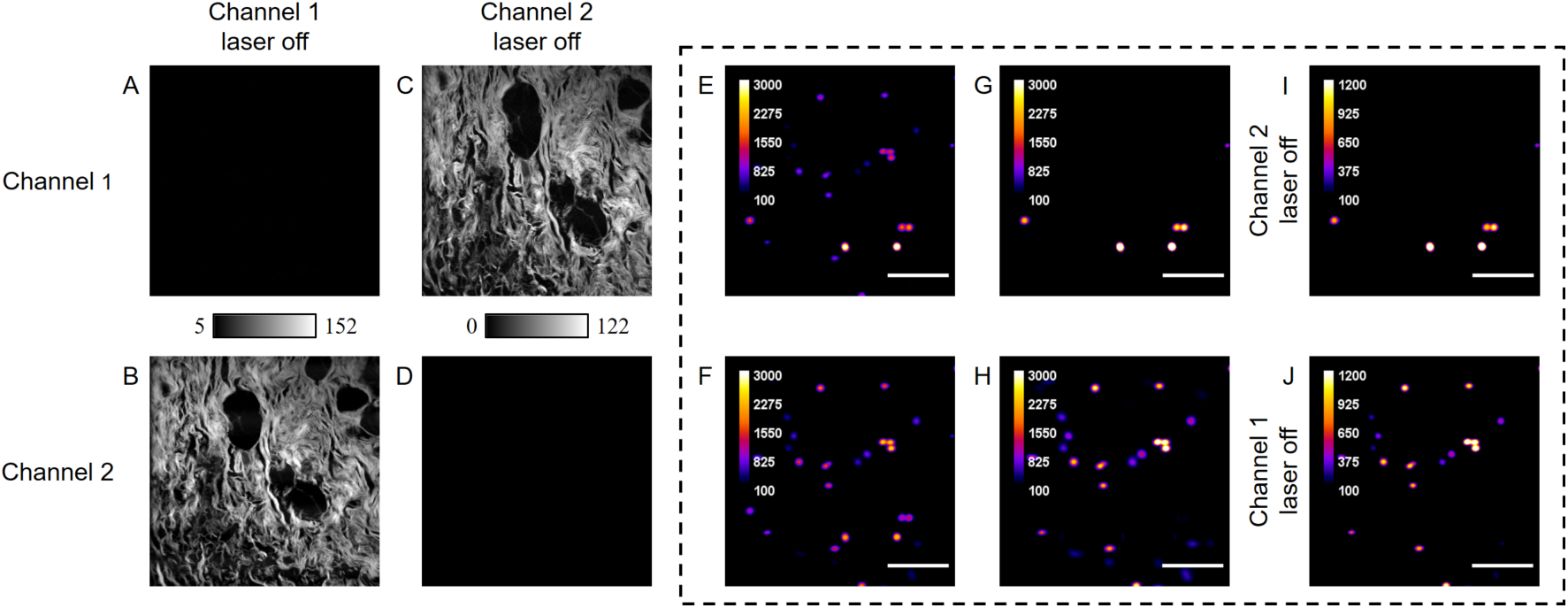
Channel cross-talk evaluation. (A). Captured Image of channel 1 when laser corresponding to channel 1 is off. (B) Captured Image of channel 2 when laser corresponding to channel 1 is off. (C) Captured Image of channel 1 when laser corresponding to channel 2 is off. (D) Captured Image of channel 2 when laser corresponding to channel 2 is off. (E and F) Averaged Image of channel 1,2 when both planes are excited. Scale bar: 20 μm. (G and H). Demixed Image of channel 1,2 when both planes are excited. (I and J). Channel 1,2 Image when laser corresponding to channel 2,1 is off.

**Fig. S2.**
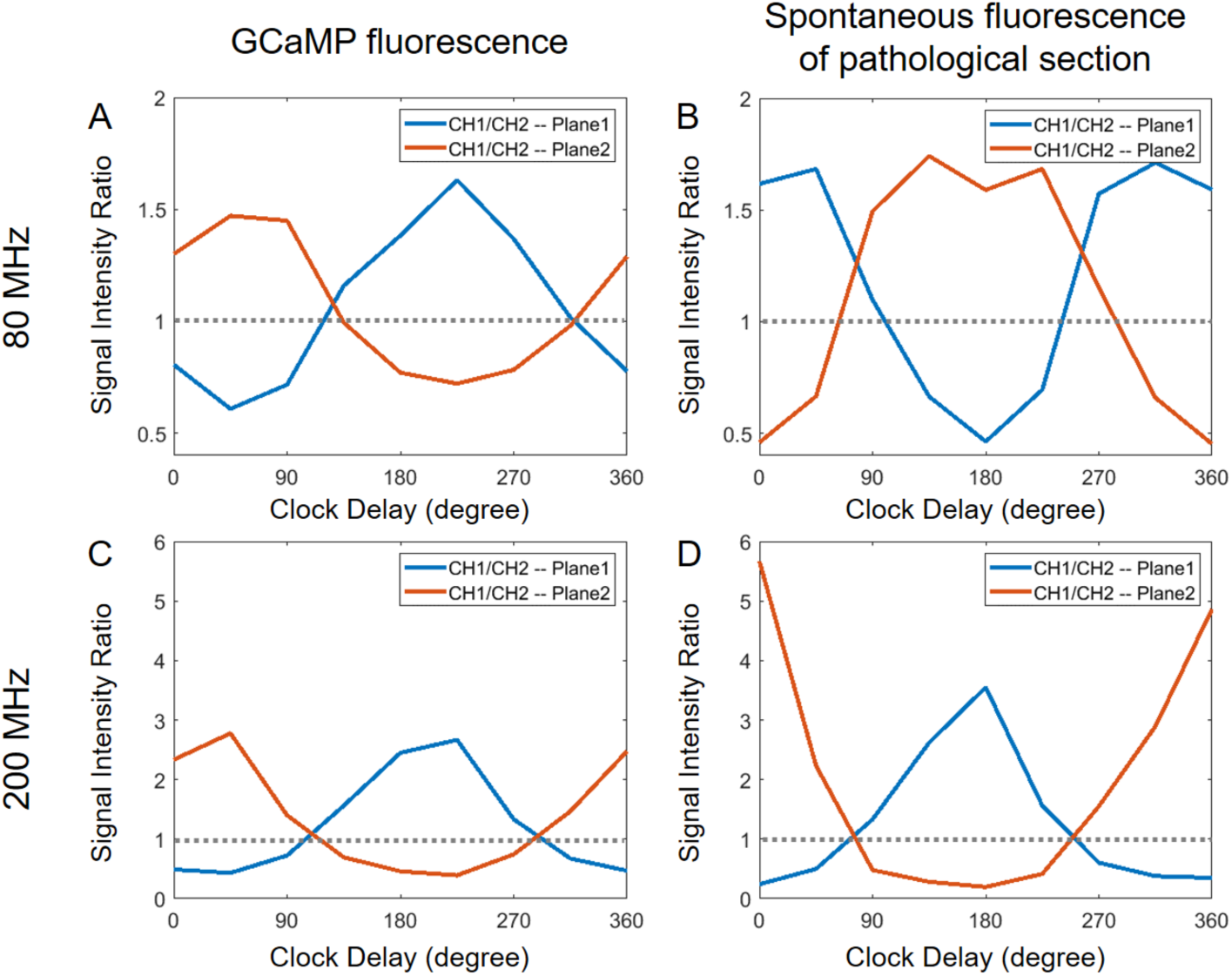
Channel crosstalk between two digitalized channels with different cut-off frequencies and biological samples. (A) Channel crosstalk with 80 MHz cut-off frequency and GCaMP labeled neuron. (B) Channel crosstalk with 80 MHz cut-off frequency and H&E stained pathological section. (C) Channel crosstalk with 200 MHz cut-off frequency and GCaMP labeled neuron. (D) Channel crosstalk with 200 MHz cut-off frequency and H&E stained pathological section.

**Fig. S3.**
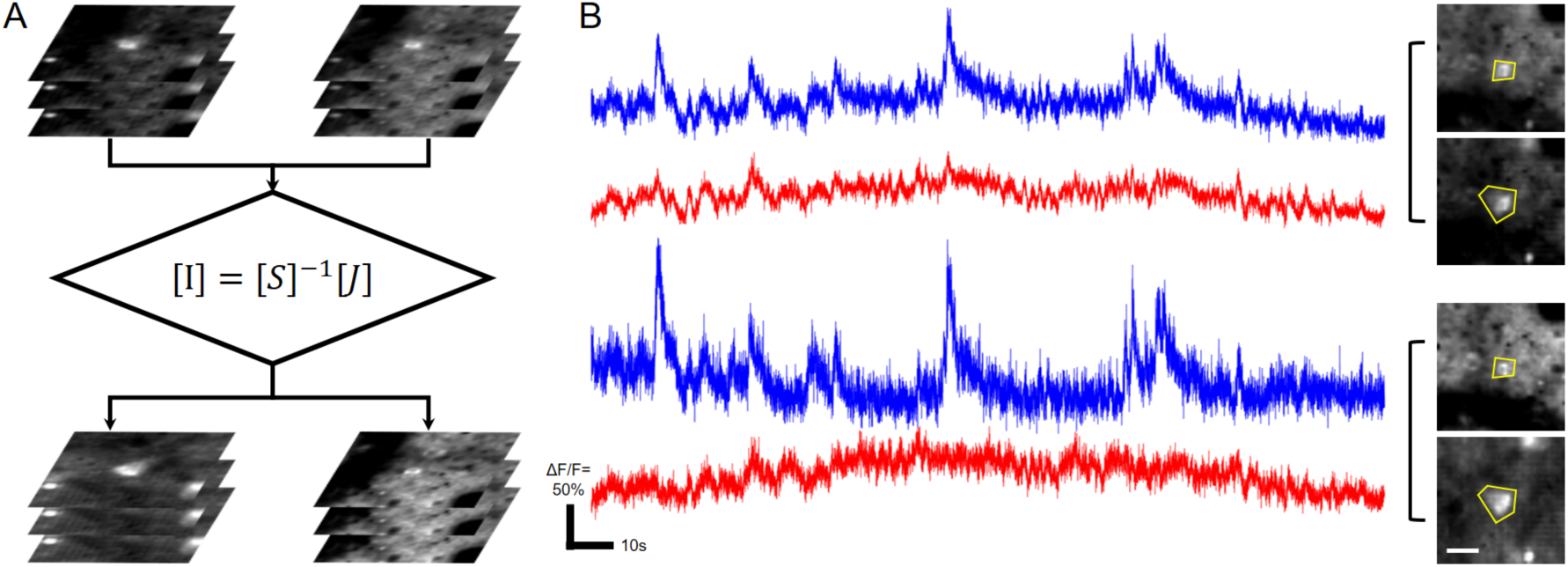
Activity traces of lateral overlap neuron from different axial depth. (A) Diagram of image linear demixing. (B) Activities of two lateral overlap neuron (blue: L2/3, red: L5) from different axial depth before(up) and after(down) linear demixing. Scale bar: 10 μm.

**Fig. S4.**
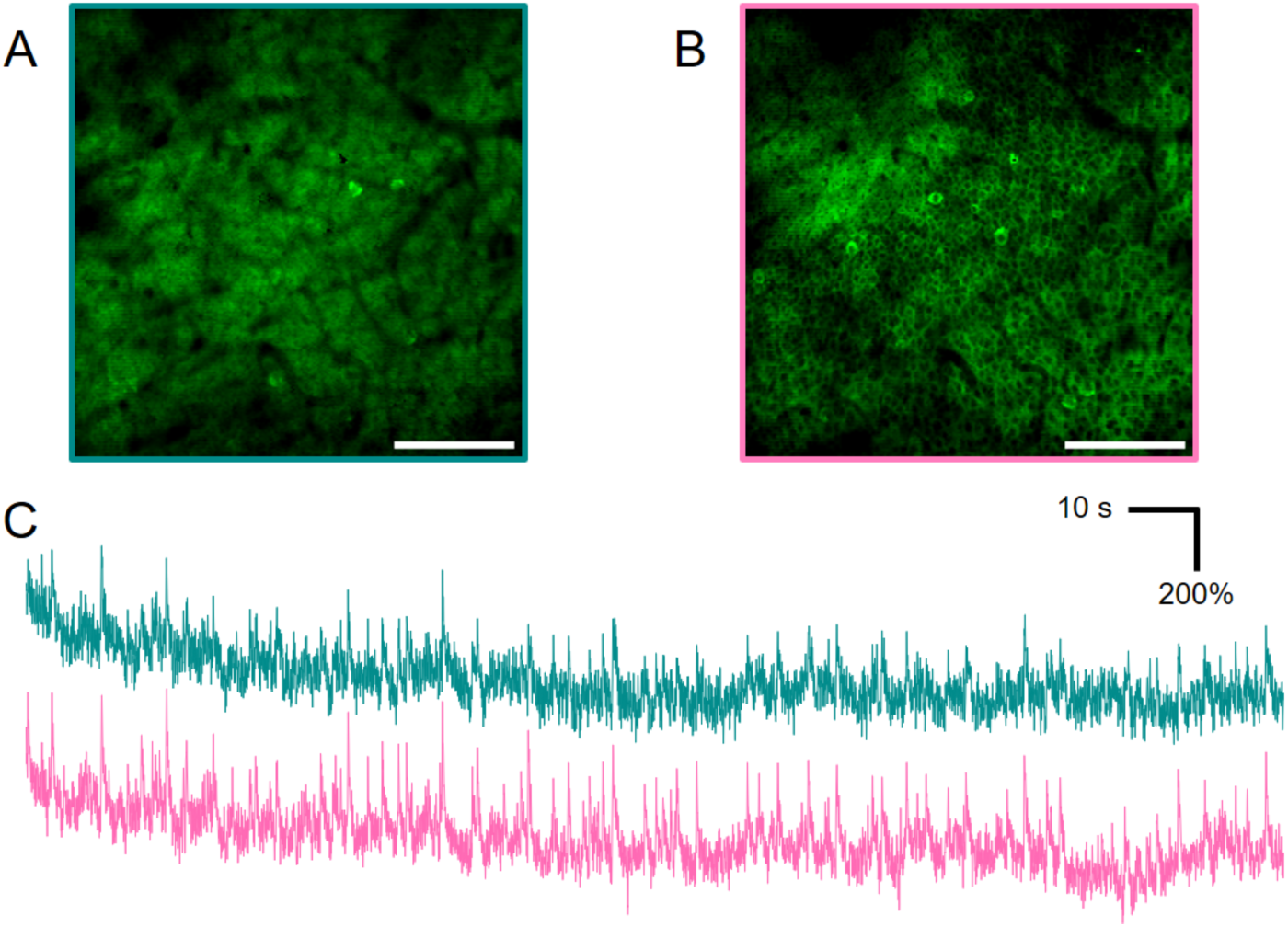
Simultaneous dual plane imaging of GABA sensor signals. (A) Averaged fields of view showing neurons expressing GABA sensor. 50 μm depth. Scale bar: 100 μm. (B) Averaged fields of view showing neurons expressing GABA sensor. 100 μm depth. Scale bar: 100 μm. (C) Z-score ΔF/F trace of GABA sensor activity from Plane 1(cyan) and Plane 2(pink). Imaging was performed on seizured mice (see Methods).

**Fig. S5.**
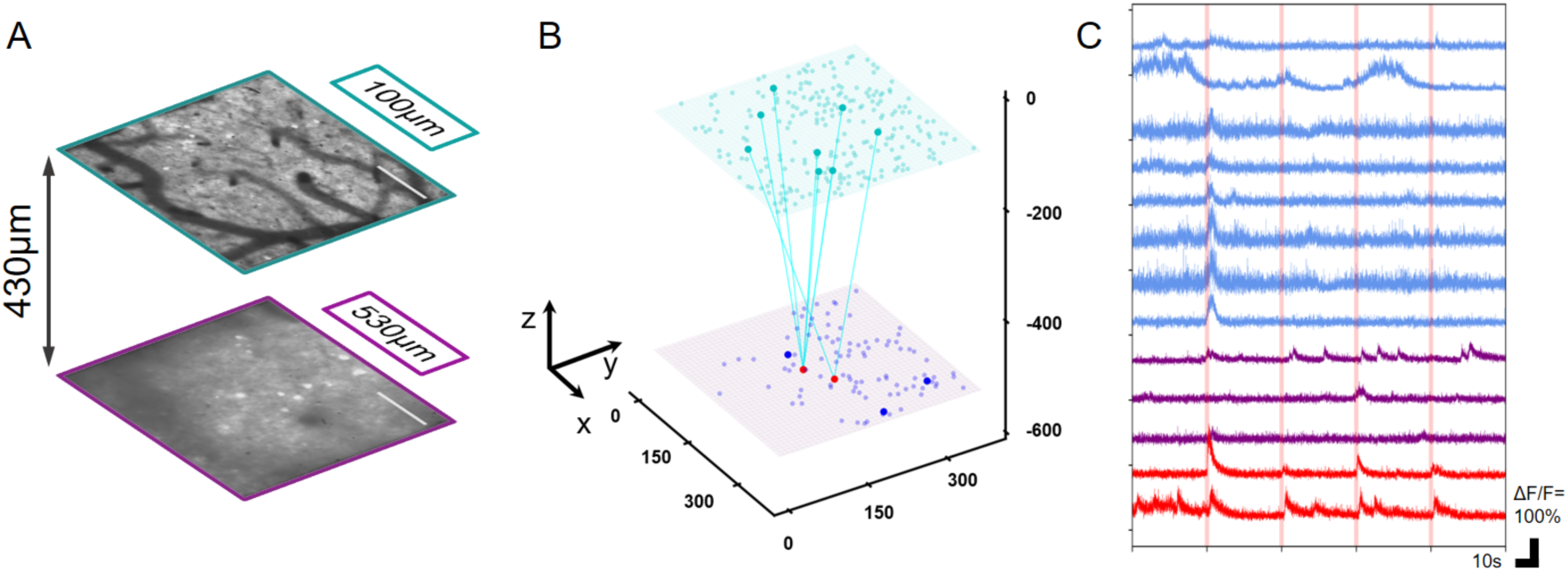
Simultaneously dual plane recording and manipulating of mouse S1 neuronal activity. (A). Averaged fields of view showing neurons expressing GCaMP6s, 100 and 530 μm depth. Scale bar: 100 μm. (B) 3D demonstration of all extracted neurons in micron. Lines between two planes indicate pairs of stimulated cells (red) and cells in response to activation (blue). Cells having response to activation in deeper plane are denoted (dark purple). (C) Traces of stimuli cells (red) in 530 μm depth and cells in response to activation in 100 μm and 530 μm depth (blue and purple).

**Fig. S6.**
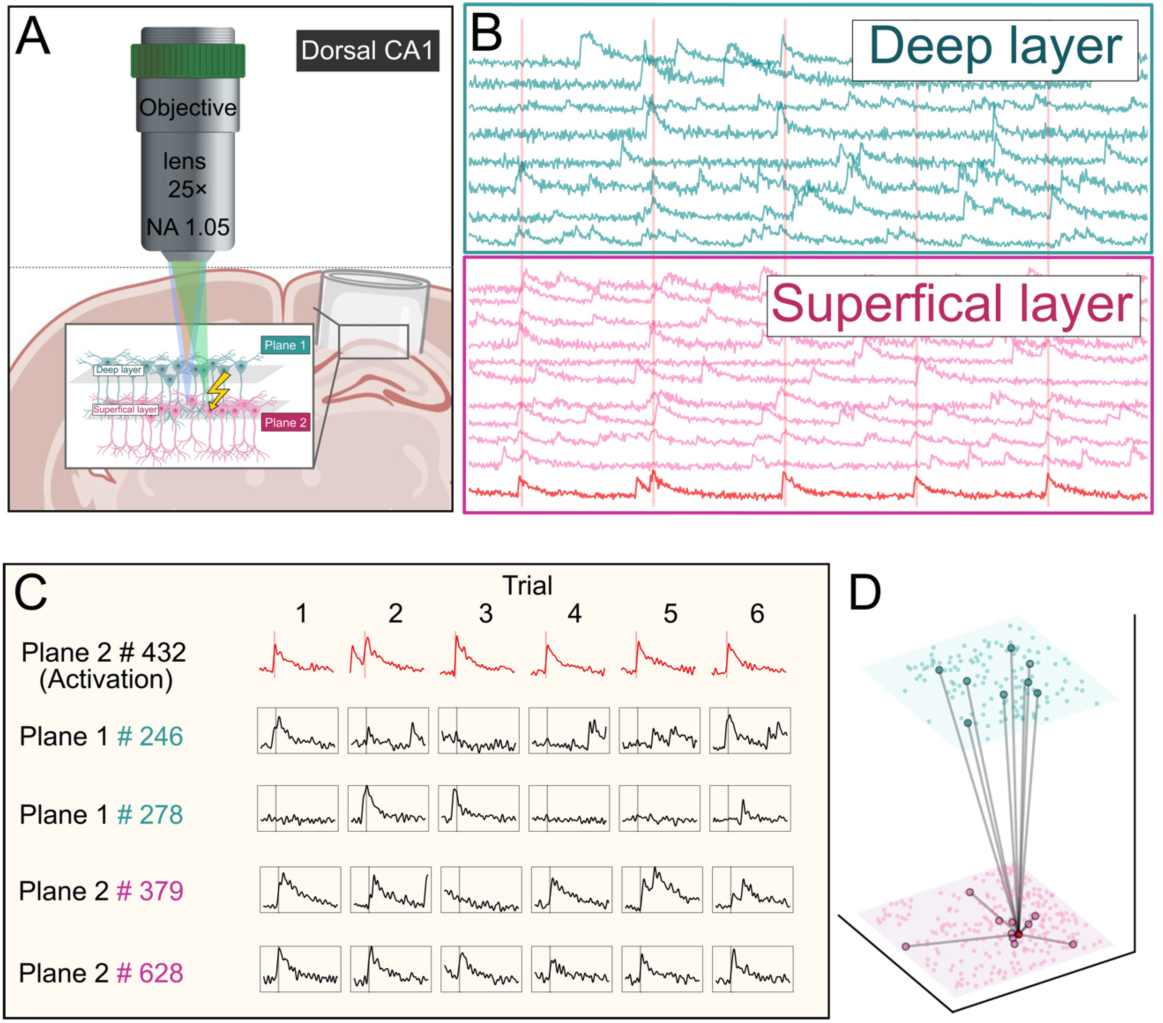
CLAOP analysis on CA1 deep and superficial layers. (A) Diagram of dual plane recording with optogenetic manipulation in mouse hippocampus CA1. (B) Activities of selected neurons from the dual plane. (cyan: upper plane, pink: lower plane, red: target cell) (C) Responses of the target neuron and other neurons to two-photon photostimulation. (D) 3D demonstration of all recorded neurons from dual-plane recording with optogenetic manipulation. Lines connecting neurons indicate a high correlation between other neurons shown in (B) and the target neuron.

**Fig. S7.**
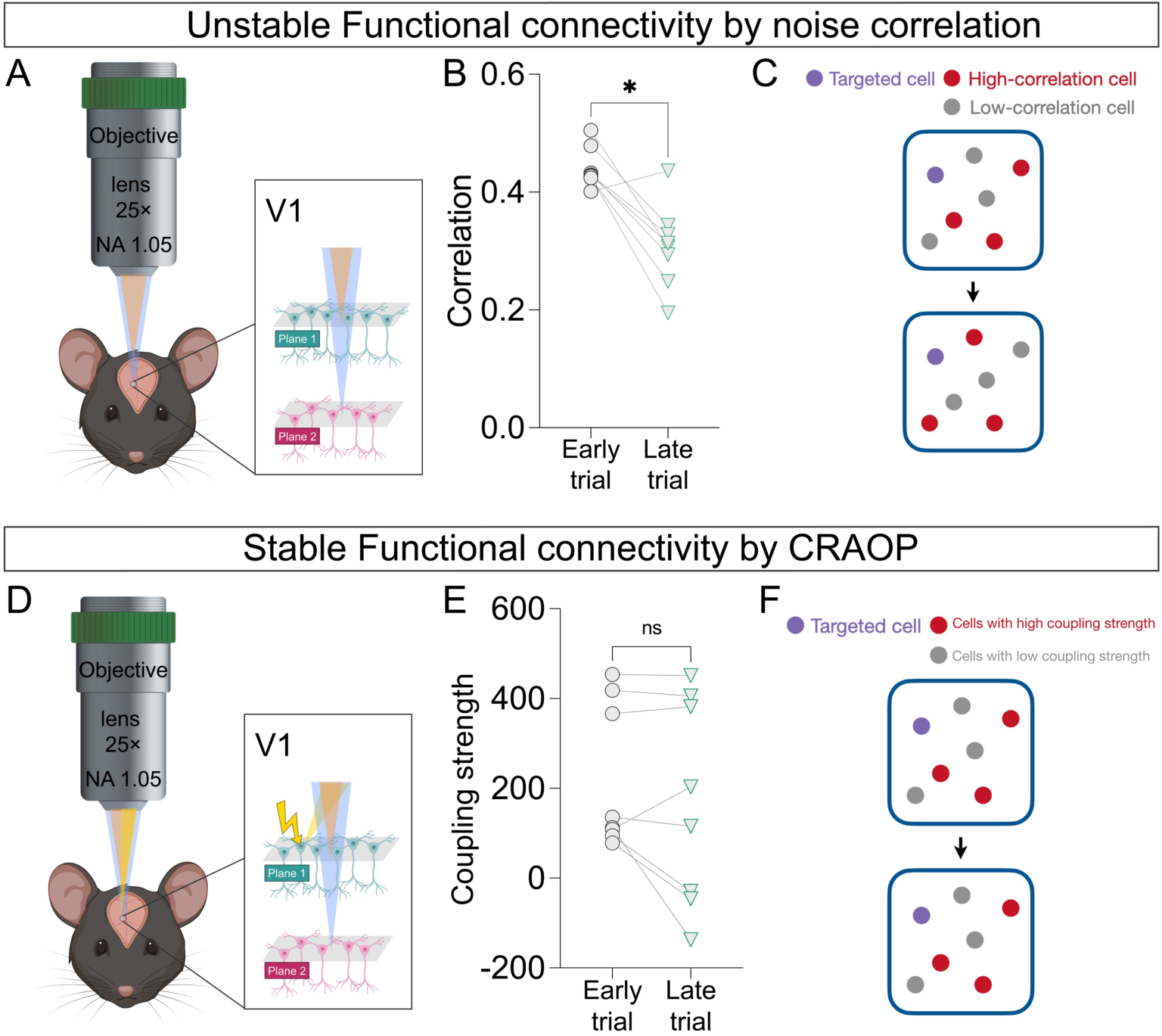
More stable functional connectivity was dissected by CLAOP. (A) Diagram of dual plane recording in mouse cortex V1. Recordings were performed on two trials where no visual stimulus was presented. (B) In the early trial, neurons with top 10% correlation in activity with a selected cell were identified. The correlations significantly decreased in the late trial. (n=8 neurons) (C) Representative diagram showing noise correlations of neurons are not stable as time elapses. (D) Diagram of dual plane recording with optogenetic manipulation in mouse cortex V1. Recordings and optogenetics stimulations on the selected cell were performed on two trials where no visual stimulus was presented. (E) In the early trial, neurons with top 10% coupling strength with the targeted cell were identified. The coupling strengths remained high in the late trial. (n=8 neurons) (F) Representative diagram showing coupling strengths dissected by CLAOP are relatively stable as time elapses. n.s. Non-significant, *P < 0.05; Two-tailed Wilcoxon signed-rank test (B and E).

**Fig. S8.**
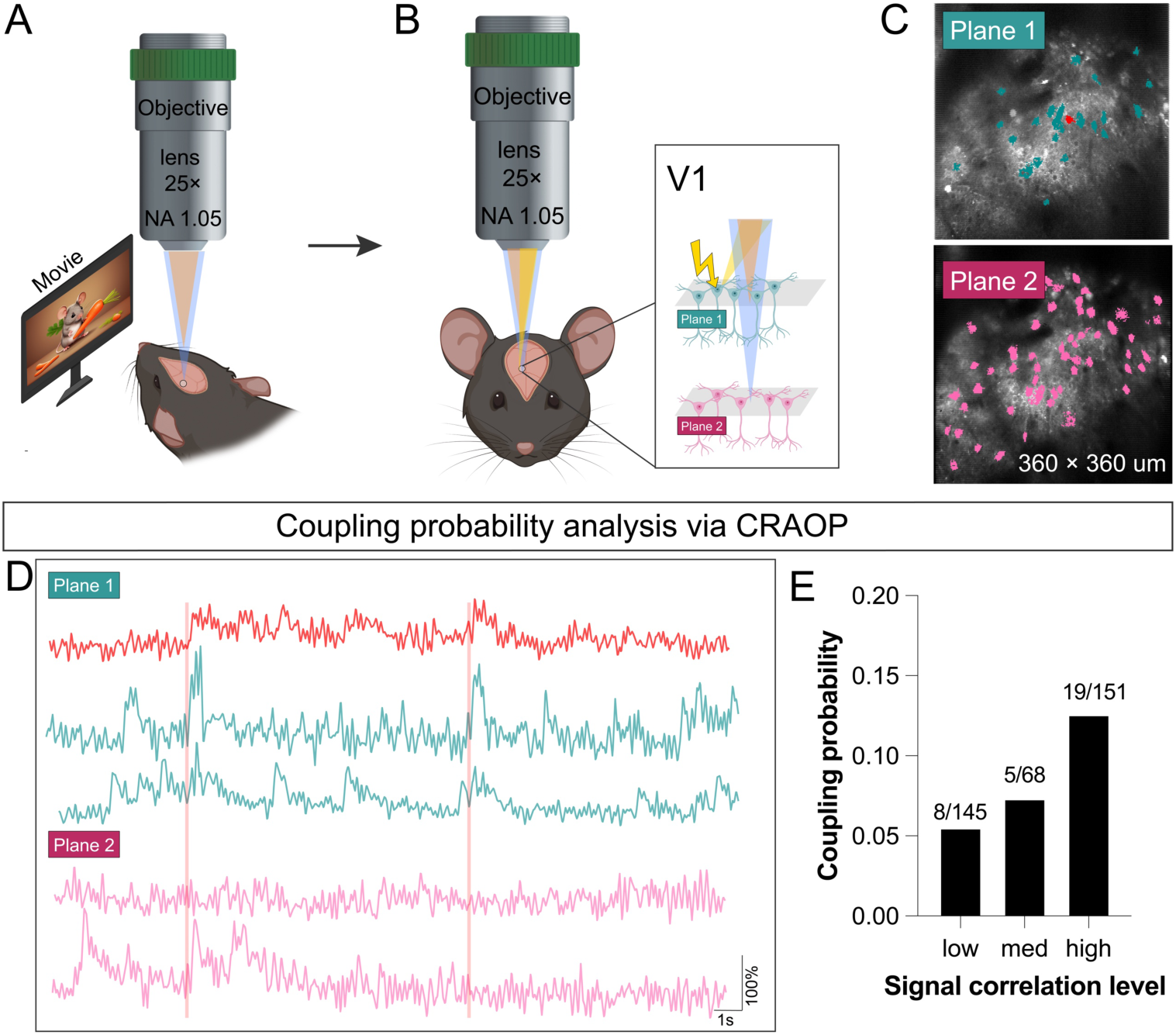
CLAOP analysis of functional connectivity with natural movies as visual stimuli correlation in V1. (A) Diagram of dual plane recording in V1 with movies as visual stimuli. (B) Diagram of dual plane recording and targeted photostimulation in V1. (C) Example of spatial masks of recorded neurons in one trial (cyan: upper plane cells, pink: lower plane cells, red: cell to be photostimulated). (D) Example of activity traces of selected neurons cyan: upper plane cells, pink: lower plane cells, red: cell to be photostimulated). Vertical lines indicate optogenetic stimulation. (E) A significant increase in coupling probability with increasing signal correlation to visual stimuli (P=0.032, Cochran–Armitage test for trend).

**Fig. S9.**
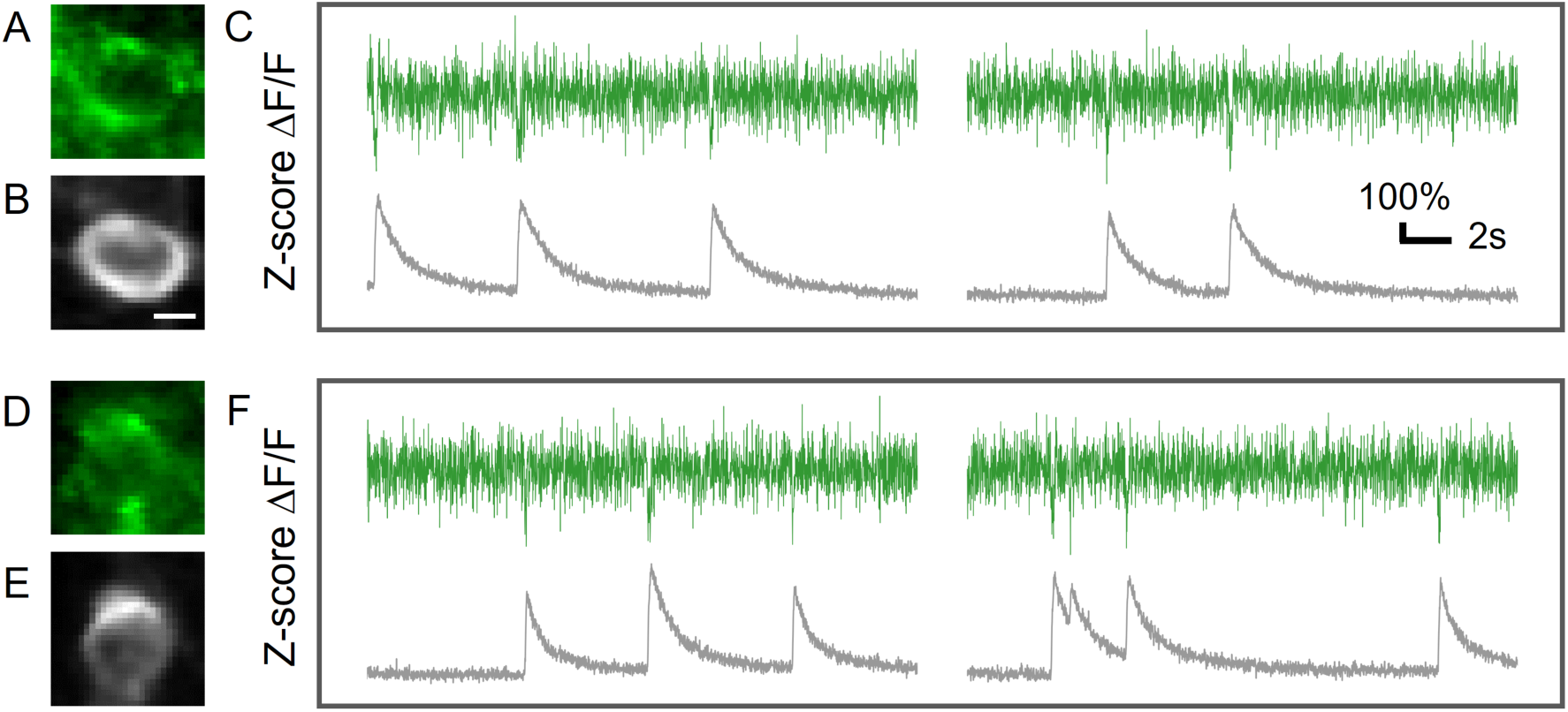
Simultaneously recording of neuronal calcium and voltage activity in mouse hippocampus CA1. (A,D) Typical neuron expressing JEDI-2P. Scale bar: 5 μm. (B,E) jRGECO expressing of the same neuron as in (A,D), respectively. (C,F) Simultaneously recording calcium and voltage activity of neurons in (A,B) and (D,E) in Z-score ΔF/F of JEDI-2P and jRGECO. Framerate: 113 Hz. The negative-going transients of JEDI-2P are in line with the onset of calcium activity peaks reflected by jRGECO.

**Table S1.** Summary of experimental conditions.

| Figure<br>s | Sample<br>Mouse<br>strain/<br>brain<br>region | Fluore<br>scent<br>label/<br>Opsin<br>label | Offset<br>lens<br>(Focal<br>length/<br>mm) | Objective | Frame<br>rate<br>(Hz) | Imaging<br>power<br>(Plane1-Plane2/<br>mW) | Power<br>used for<br>photosti<br>mulation | photostim<br>ulation:<br>reps ×<br>stimulatio<br>n_duration |
| --- | --- | --- | --- | --- | --- | --- | --- | --- |
| 2(a-e) | Ai162/V<br>1 cortex | GCaM<br>P6s | 75 | CFI75<br>LWD 16X<br>W | 60 | 60-179 |  |  |
| 2(f-j) | B6/CA1 | GCaM<br>P8m | 250 | XLPLN25X<br>SVMP2 | 60 | 24-24 |  |  |
| 2(k-n) | Ai162/<br>S1<br>cortex | GCaM<br>P6s | 250 | XLPLN25X<br>SVMP2 | 396.1 | 22-26 |  |  |
| 2(o-s) | Ai162/S<br>1 cortex | GCaM<br>P6s | 75 | CFI75<br>LWD 16X<br>W | 30/10 | 75-195 |  |  |
| 3(a-d) | Thy1-GCaMP<br>6s/CA3 | GCaM<br>P6s<br>/ChR<br>mine | 250 | XLPLN25X<br>SVMP2 | 60 | 43-108 | 128 mW | 100×1 ms |
| 3(e-j) | B6/CA1 | JEDI2<br>p/C1V<br>1 | 250 | XLPLN25X<br>SVMP2 | 396.1 | 88-107 | 128mW | 100×10<br>ms |
| 3(i-n) | Ai162/S<br>1 cortex | GCaM<br>P6s/C<br>1V1 | 75 | CFI75<br>LWD 16X<br>W | 30/10 | 75-195 | 153mW | 100×10<br>ms |
| 4(a-d) | B6/ V1<br>cortex | GCaM<br>P8m/rs<br>ChRm<br>ine | 250 | XLPLN25X<br>WMP2 | 60 | 56-104 |  |  |
| 4(e-g) | B6/ V1<br>cortex | GCaM<br>P8m/rs<br>ChRm<br>ine | 250 | XLPLN25X<br>WMP2 | 60 | 56-104 | 170mW | 100×4 ms |
| 5(a-g) | Ai162/<br>V1<br>cortex | GCaM<br>P6s/C<br>1V1 | 250 | XLPLN25X<br>WMP2 | 60 | 42-63 |  |  |
| 5(h-n) | Ai162/<br>V1<br>cortex | GCaM<br>P6s/C<br>1V1 | 75 | CFI75<br>LWD 16X<br>W | 30 | 28-137 | 153mW | 100×5 ms |
| S4 | /CA1 | GABA sensor | 250 | CFI75 LWD 16X W | 60 | 45-45 |  |  |
| S5 | Ai162/S1 cortex | GCaM P6s | 75 | CFI75 LWD 16X W | 60 | 60-179 | 153mW | 100×10 ms |
| S6 | B6/CA1 | GCaM P8m | 250 | XLPLN25X SVMP2 | 60 | 24-24 |  |  |
| S7 | B6/ V1 cortex | GCaM P8m/rs ChRmine | 250 | XLPLN25X WMP2 | 60 | 56-104 | 170mW | 100×3 ms |

**Movie S1 (separate file).** High frame rate (396.1Hz) dual plane imaging of GCaMP6s-labeled neurons in layer2/3 of the mouse sensory cortex. The pixel number of this video is 256 x 24.

**Movie S2 (separate file).** Dual plane recording with optogenetic manipulation in mouse hippocampus CA1. We conducted continuous imaging with 9 trials of stimulus. The pixel number of this video is 256 x 256.

## Appendix

For temporal multiplexing imaging of N planes, detected signal [J] can be described as following equations:

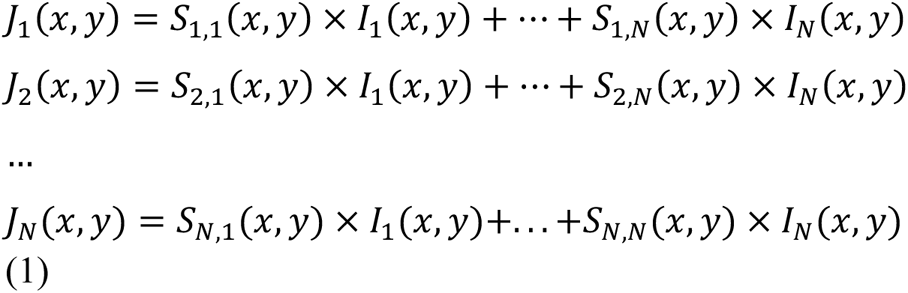

Here, *S_i,j_* is channel crosstalk from plane *j* to channel *i*, and following equation holds:

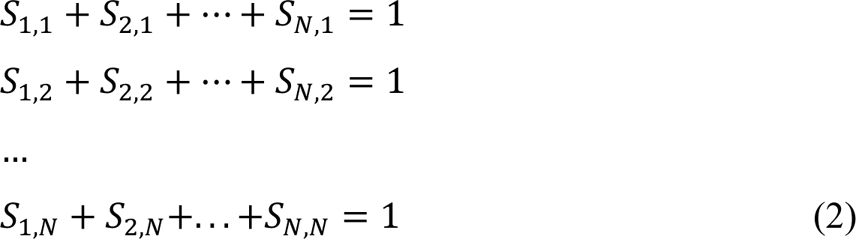

Thus, original signal [I] from different planes can be calculated as:

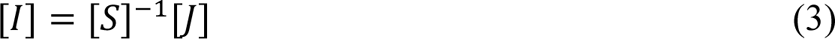

For our dual plane imaging configuration, detected signal [J] of each channel can be described as following equations:

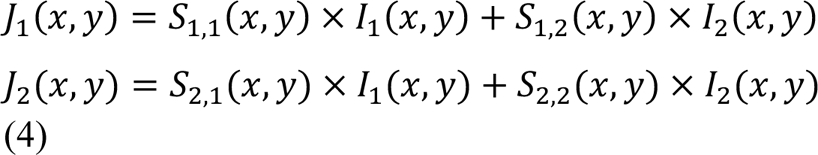

Elements of crosstalk [S] can be determined based on individually imaging results of lower plane and upper plane. It can also be determined by looking for a coefficient that minimizes the correlation of the demodulated images. The matrix [S] changes with different indicators and varying bandwidth of transimpedance amplifier. In our experiment, lower cut-off frequency of transimpedance amplifier leads to higher SNR of imaging results. However, high signal aliasing will also increase the amplitude of additive noise.

Considering the actual situation with noise *N_i_* on each channel:

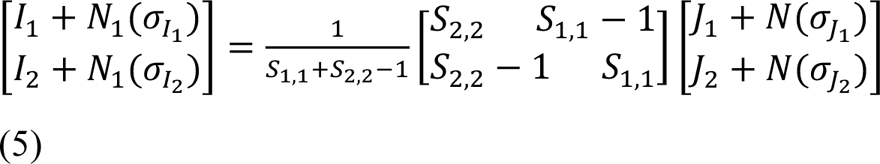

When S_1,1_ and S_2,2_ are nearly 1, it is close to the ideal situation without signal aliasing. The amplification effect of noise is negligible. However, when S_1,1_ and S_2,2_ are approaching 0.5 (totally aliasing), noise of recovered signal may be amplified gradually. Taking plane 1 as an example:

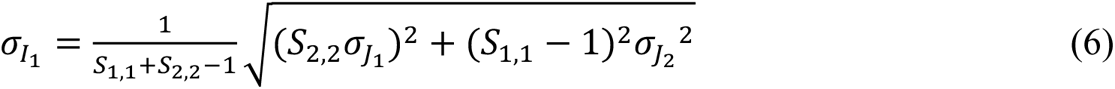

Assuming S_1,1_ = S_2,2_ = 0.7 and *σ*_J1_ = *σ*_J2_ = *σ*, we get *σ*_I1_= 1.90 *σ*, which means that noise is amplified compared to aliased raw image. Adopting a data acquisition board with a high sampling rate (up to GHz) or reducing the repetition rate of the femto-second laser may reduce channel crosstalk and improve the performance of temporal multiplex imaging.

To demonstrate that our system can realize dual-plane neural activity recording without the need for image sparsity or prior knowledge. We select two neurons that have lateral overlap but from different axial plane (100 μm and 530 μm). The raw data is first redistributed by linear demixing (**fig. S3A**). Averaged images of two neurons and the corresponding calcium signal are shown in **fig. S3B**. It is shown that neuron activities of two lateral overlap neurons can be extracted distinctly. The correlation coefficient of traces from raw data is 0.8177, which indicates high aliasing before linear demixing. However, the correlation coefficient of traces is reduced to -0.0904, which shows that the calcium signal of two neurons can be reliably recorded.

